# Ecological inheritance facilitates the coexistence of environmental helpers and free-riders

**DOI:** 10.1101/2025.06.30.662428

**Authors:** Iris Prigent, Charles Mullon

## Abstract

Variation in social traits and behaviours is widespread in nature and can be maintained by selection. Many social traits influence fitness indirectly by modifying shared environments that are transmitted across generations, a process known as ecological inheritance. Here, we investigate how variation in environmentally mediated social behaviour, such as helping to improve shared resources, can emerge and persist in spatially subdivided populations where locally modified environments are transmitted across generations. Using mathematical and computational modelling, we show that ecological inheritance, when combined with limited dispersal, readily leads to the stable coexistence of two types with opposite environmental legacies: helpers, who improve the local environment for the future at a personal cost, and environmental free-riders, who benefit without contributing to the detriment of future generations. In turn, this polymorphism generates lasting spatial heterogeneity in environmental quality and, consequently, in survival and reproduction—particularly under isolation-by-distance, creating stable clusters of high- and low-quality habitats across an otherwise homogeneous landscape. These findings reveal how ecological inheritance and spatial structure interact to stabilise polymorphism, potentially driving long-term behavioural, ecological, and fitness variation across diverse biological systems.

## 1 Introduction

Within virtually all populations, individuals vary in the way they interact with conspecifics (e.g. in their tendencies to help, Trivers, 1985; Crespi, 2001; Koenig and Dickinson, 2004; Komdeur, 2006; compete with, Skulason and Smith, 1995; Smith and Skúlason, 1996; Bolnick et al., 2003, 2011; or punish others, Trivers, 1985; Clutton-Brock and Parker, 1995; Riehl and Frederickson, 2016). Adaptive variation in social behaviour persists over time if different genetic variants associated with variation in these behaviours can stably coexist, a scenario consistent with frequency-dependent selection (whereby the fitness of a variant changes with its frequency, e.g. Wright, 1948; Lewontin, 1958; Maynard Smith, 1962; Clarke and O’Donald, 1964; Ayala and Campbell, 1974; Maynard Smith, 1982; Heino et al., 1998; Leimar and McNamara, 2023; Ruzicka et al., 2025; Lehmann and Mullon, 2025). This can happen through a wide range of social interactions, either direct – such as combat (e.g. Cavalli-Sforza and Feldman, 1978; Maynard Smith, 1982; Doebeli and Hauert, 2005) – or indirect, mediated through the environment – such as the consumption of shared resources or the production of a common good (e.g. Slatkin, 1980; Doebeli and Dieckmann, 2000; Ajar, 2003; Rueffler et al., 2006; Brown and Taddei, 2007; Kisdi and Geritz, 2010; Wakano and Lehmann, 2014; Schmid et al., 2024, for recent review, Lehmann and Mullon, 2025).

When environmental effects carry over across generations, a phenomenon known as ecological inheritance (Odling-Smee, 1988; Odling-Smee et al., 2003; Pontarotti, 2022), understanding the evolutionary emergence and maintenance of adaptive variation becomes challenging as the fitness of a trait variant depends not only on its current frequency but also on its past frequency. Several models have investigated this, typically assuming that the environmental effect of trait expression by an individual is experienced by the entire population (i.e. in well-mixed populations, Odling-Smee et al., 2003; Brown and Taddei, 2007; Weitz et al., 2016; Tilman et al., 2020; Ito and Yamamichi, 2024). For instance, Weitz et al. (2016) studied the coupled dynamics of environmental quality and the frequency of two types: “cooperators” who improve the environment and “defectors” who deteriorate it. These two types can stably co-exist when defectors have a fitness advantage in a good environment, while cooperators have an advantage in a bad one. By generating a time-lag between type frequency and environmental consequences, ecological inheritance destabilises coexistence, generating eco-evolutionary cycles alternating between two extreme states, one dominated by cooperators and the other by defectors. Similar destabilizing effects have been observed in other models assuming well-mixed populations (Brown and Taddei, 2007; Tilman et al., 2020; Ito and Yamamichi, 2024).

Given the physical limitations of movement and interactions (Clobert, 2012), most natural populations are in fact dispersal-limited and the environmental effects of a trait limited to the vicinity of the individuals expressing it, i.e. effects are local. How this spatial structure influences adaptive trait variation under ecological inheritance is not straightforward, as selection depends on current and past local frequency of traits that fluctuate in time due to local sampling effects under limited dispersal. Due to such sampling effects, the genes underlying a trait and the environmental effects of this trait become statistically associated, with individuals preferentially inheriting environments modified by their ancestors. From the perspective of these genes, the local environment becomes part of the “extended phenotype” (Dawkins, 1982, 2004), and selection on traits due to their environmental effects is influenced by kin selection (Lehmann, 2008). Current analyses of selection on traits under ecological inheritance and limited dispersal mostly focus on the direct effects of traits on invasion fitness (i.e. how a trait change in a mutant influences its own growth rate, e.g. Rousset and Ronce, 2004; Lehmann, 2007, 2008; Sozou, 2009; Mullon and Lehmann, 2018; Mullon et al., 2021, 2024). However the emergence of adaptive polymorphism depends on how the growth rate of a rare mutant changes when the trait values in the resident population also change (Geritz et al., 1998; Lehmann and Mullon, 2025), and only recently have theoretical advances made analysis of this phenomenon possible under ecological inheritance and limited dispersal (Ohtsuki et al., 2020; Prigent and Mullon, 2023). Here, we leverage this recent theory to explore the evolution and ecological consequences of a trait with positive long-term and local environmental effects in a subdivided population with limited dispersal, with specific interest in the conditions under which polymorphism emerges and is maintained.

## 2 Model

### 2.1 Biological scenario

We consider a population subdivided among a large number of patches. Each patch is characterised by an environmental variable *ϵ* ∈ ℝ that is constructed or engineered by the individuals residing in it and that increases their fitness. This variable represents a good that is accessible to every individual in the patch, e.g. the amount of reserves in transformed food, the quality of built nesting sites, or the concentration of an excreted chemical compound. Individuals express a quantitative trait *z* ∈ ℝ that is individually costly but increases the environmental variable of their patch, e.g. the effort in constituting and maintaining reserves of degradable resources, in building structures to protect against biotic or abiotic attacks, or in excreting useful chemicals that are otherwise unavailable. More broadly, *z* should be thought of as the costly investment in a local good that is transferable across generations. As such, the trait *z* represents a form of helping that occurs via the external environment, which exhibits ecological inheritance.

We are interested in the effects of ecological inheritance on whether selection favours polymorphism in helping (i.e. in trait value *z*), and in turn, how polymorphism influences heterogeneity in the environment (i.e. in the environmental variable *ϵ*). To focus on ecological inheritance, we assume that generations do not overlap (using a Wright-Fisher model of reproduction) and that individuals are haploid, and to focus on the effects of polymorphism on environmental heterogeneity, we assume that in the absence of variation in *z*, the environmental variable stabilises to the same value in all patches (i.e. the environment is homogeneous).

Individuals go through the following life-cycle (Fig. 1A): (i) adults interact with one another and with their local environment, transforming the environmental variable of their patch; (ii) adults reproduce clonally, with a fecundity that depends on these interactions; (iii) each offspring either remains in its natal patch (with probability 1 − *m*) or disperses (with probability *m*); (iv) adults die and offspring compete locally to become the adults of the next generation. We initially assume that patches are arranged according the island model of dispersal (i.e. there is no isolation-by-distance so when an offspring disperses, it is equally likely to land in any other patch), and that all patches carry a fixed number *N* of adults. We relax both of these assumptions later.

**Figure 1:**
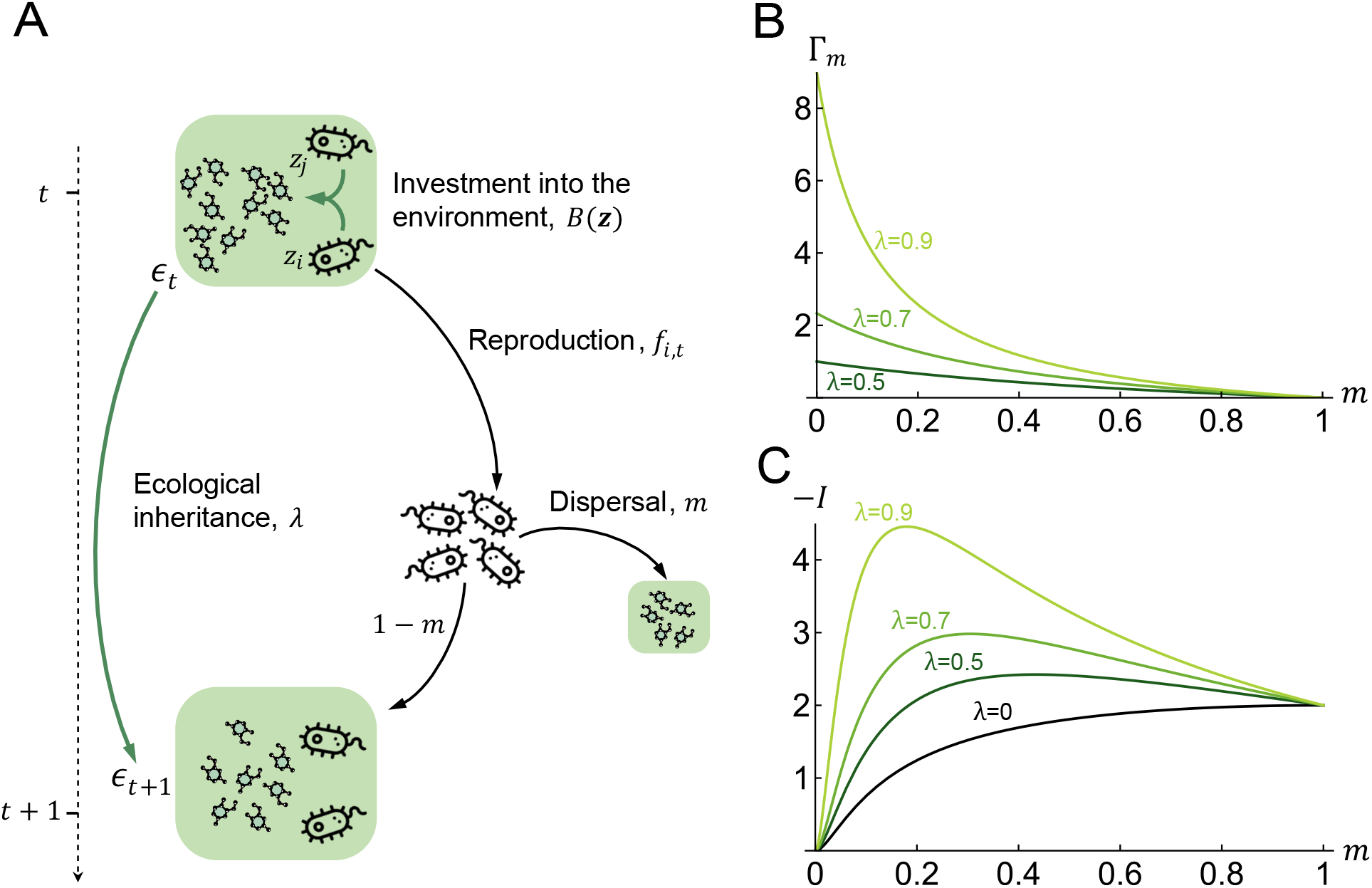
**(A) A model of inter-generational helping via environmental modifications**. Individuals – represented as free-living cells – reside in groups characterized by an environmental variable *ϵ*_*t*_, e.g. the density of fitness-enhancing extracellular molecules as shown. Each individual invests some effort *z* that increases *ϵ*_*t*_, of which a proportion *λ* is transmitted to the next generation due to ecological inheritance (eq. 1). Individuals reproduce with fecundity that decreases with their investment *z* but increases with the environmental state of their patch *ϵ*_*t*_ (fecundity given by eq. 4). Offspring then either disperse with probability *m* or remain in their patch with probability 1 −*m*, in which case they get to experience the environmental legacy of their parent (when *λ >* 0). See section 2.1 for more details on the model. **(B) Strength of selection on inter-generational helping, Γ**_***m***_. Selection for *z* to evolve is strongest when dispersal *m* is weak and ecological inheritance *λ* is strong (from eq. 7, Lehmann, 2007; section 2.3 here). **(C) Potential for coexistence between environmental helpers and free-riders** −***I***, which determines whether two types close to the equilibrium *z*^*^ can mutually invade one another and therefore be maintained (from eq. 10; other parameters: *N =* 10, *b*_3_ *=* −2.5.). This shows coexistence is more stable for ecological inheritance *λ* and intermediate dispersal *m*. Section 3.1 for interpretation.

### 2.2 Environmental transformations and individual fitness

To specify the inter-generational transformation of the environment through trait expression, consider an arbitrary generation *t* and let *ϵ*_*t*_ denote the environmental variable of a patch at the start of generation *t* (i.e. before step i of the life-cycle). Let ***z***_*t*_ *=* (*z*_1,*t*_, …, *z*_*N,t*_) be the collection of traits expressed by the adults indexed 1 to *N* in that same patch (where *z*_*i,t*_ is the trait of adult indexed *i* ∈ {1,…, *N*}). We assume that the environmental variable of the patch at the next generation is

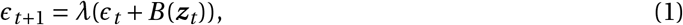

where the parameter 0 ≤ *λ <* 1 tunes the degree of ecological inheritance, and the function 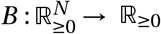 determines the effects of trait expression on the environmental variable. Eq. (1) states that individuals at generation *t* start with environmental variable *ϵ*_*t*_, to which they add *B* (***z***_*t*_), and finally a proportion *λ* of this total is transmitted to the next generation (we consider more general functions in our appendix and refer to these results in our discussion).

To model the influence of individuals on their local environment, we decompose the function *B* as,

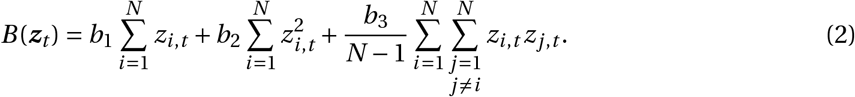

The parameters *b*_1_ and *b*_2_ determine the marginal effect an individual has on the environment, with *b*_1_ giving the additive (or first-order) effect and *b*_2_ the non-additive (or second-order) effect. We assume that *b*_1_ *>* 0 such that in the absence of helping (i.e. when *z =* 0), a trait increase in one individual has a positive marginal effect on the environment. As it increases, this individual effect may eventually saturate (when *b*_2_ *<* 0) or amplify (when *b*_2_ *>* 0). The parameter *b*_3_ meanwhile captures the effect of interactions among individuals on the environment: when *b*_3_ *<* 0, the interaction is negative or antagonistic, i.e. the increase in trait value by one unit in two individuals has a lesser effect on the environment than the increase in trait value by two units in one individual; Conversely, the interaction is positive or synergistic when *b*_3_ *>* 0, so that the increase in trait value by one unit in two individuals has a greater effect on the environment than the increase in trait value by two units in one individual. To ensure that *B* (*z*) always increases with an increase in trait value in one individual, i.e. that *z* always constitutes a form of indirect helping, we must have that

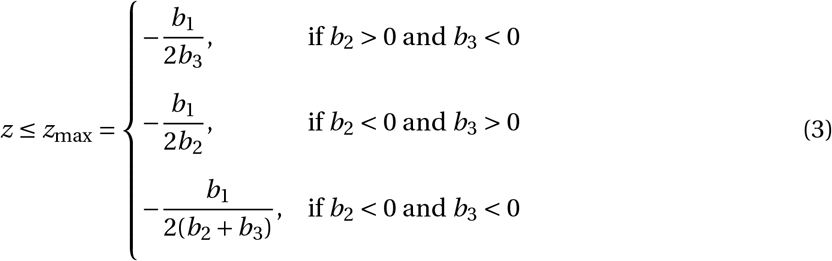

(Appendix A.1 for details).

The fecundity of an individual depends on the balance between on one hand, the costs of expressing trait value *z*, and on the other hand, the benefits of residing in a patch that shows a “good” environment *ϵ*. Specifically, we let the fecundity of a focal individual *i* at generation *t* be

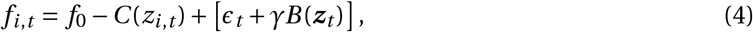

where *f*_0_ is baseline fecundity; *C* : ℝ_≥0_ → ℝ_≥0_ is a cost function that depends on the trait *z*_*i,t*_ of the focal,

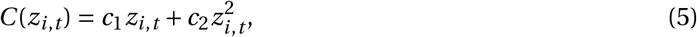

with *c*_1_ and *c*_2_ as parameters that respectively determine the additive and non-additive marginal effects of trait expression on fecundity. Finally, the term within square brackets in eq. (4) can be interpreted as follows: individuals of generation *t* in the focal patch began with *ϵ*_*t*_, to which they added *B* (***z***_*t*_), and a proportion 0 ≤ *γ* ≤ 1 of their own contribution they get to experience in their lifetime. The parameter *γ* thus tunes the amount of direct benefit that an individual enjoys from its own contribution to the environment.

### 2.3 Evolution of helping through the local environment

The forms of eqs. (1)-(5) allow us to connect to several models of social evolution, notably from classical evolutionary game and niche construction theory (Appendix B for details). The most relevant connection is with Lehmann (2007), which our baseline model falls under. Lehmann (2007) (his eq. 9) shows that in this model, selection favours the emergence of helping through the environment, i.e. an increase in *z* in a population where *z =* 0, when

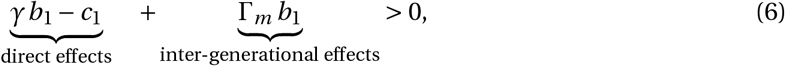

where

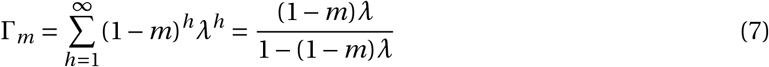

is the environmental effect that an individual has on all its descendants (on average, (1 −*m*) descendants that remain in their patch and thus inherit an environmental effect *λ* at each generation; see Appendix C.3 here for a re-derivation of eq. 6). Inequality (6) shows that in the absence of ecological inheritance (*λ =* 0), helping only evolves if an individual receives more direct benefits (*γb*_1_) from increasing its trait value than the cost of this increase (*c*_1_) (Taylor, 1992). Ecological inheritance (*λ >* 0) boosts positive selection for helping as by increasing its trait value and improving its patch, an individual now also benefits its descendants who live in that patch (Fig. 1B).

Inequality (6) gives the condition for the evolution of helping through the local environment. Once helping has evolved, are there conditions under which helping experiences negative frequency-dependent selection such that individuals showing different levels of helping coexist? We investigate these conditions under the assumptions that trait *z* evolves under the constant input of rare mutations with small phenotypic effect. Our analyses can be found in Appendices C-G. Our main results are summarized below.

## 3 Results

### 3.1 Ecological inheritance favours mutual invasion and thus coexistence

Once it has emerged, we show that helping evolves to the evolutionary equilibrium (i.e. the convergence stable trait value)

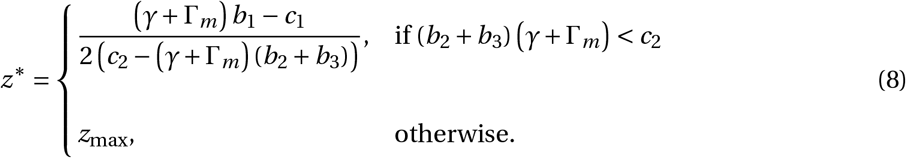

(Appendix C.3 for derivation). Unsurprisingly, greater levels of helping evolve when its benefits are large and its costs are small (i.e. *z*^*^ increases with *b*_1_ and *b*_2_ *+ b*_3_, and decreases with *c*_1_ and *c*_2_). Additionally, *z*^*^ increases with *γ* and Γ_*m*_ (Supp. Fig. S1); that is, *z*^*^ is greatest where the recipients that benefit from an individual investing into the environment are: itself (large *γ*) and its descendants (large Γ_*m*_). In a population that has converged to express *z*^*^, all patches then show the same ecological equilibrium

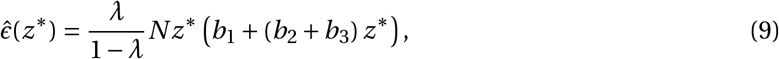

obtained from solving 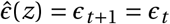 (using eq. 1) at *z*^*^. This ecological equilibrium 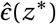 increases with ecological inheritance *λ* as greater *λ* causes selection to favour greater investment *z*^*^ into the environment (eq. 8), and also allows for greater inter-generational accumulation of environmental benefits (an accumulation which becomes extremely large in eq. (9) as *λ* approaches one).

To investigate the conditions that favour variation in trait *z*, we first study when selection favours the the stable coexistence of two types of individuals: (i) some that invest more into the environment than *z*^*^ (expressing trait value *z*_*+*_ *= z*^*^ *+* Δ); and (ii) some that invest less (expressing trait value *z*_−_ *= z*^*^ − Δ). We show in Appendix C.5 that when Δ is small, the reciprocal invasion of *z*_*+*_ by *z*_−_ and of *z*_−_ by *z*_*+*_ is determined by

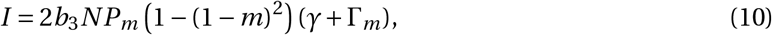

where

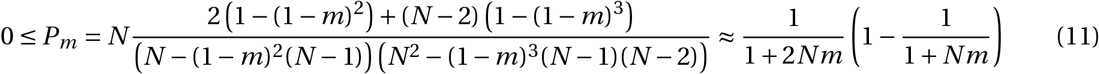

(the approximation for *P*_*m*_ is obtained by taking the limit *m* → 0 with *Nm* constant, e.g. Wright, 1931). The quantity *I* (eq. 10) must be negative for *z*_*+*_ and *z*_−_ to reciprocally invade one another and therefore for these two types to coexist (specifically, −*I* is indicative of the growth rate of *z*_*+*_ when it is rare in a population where *z*_−_ is common and of *z*_−_ when it is rare in a population where *z*_*+*_ is common).

Eq. (10) reveals that whether coexistence is possible is determined by the sign of *b*_3_ (as the rest of eq. 10 is greater or equal to zero). Specifically, coexistence is possible when *b*_3_ *<* 0, that is when interactions among individuals have antagonistic effects on the environmental variable of their patch (recall eq. 2). This occurs when the environmental benefits *B* (***z***) saturates with the number of *z*_*+*_ individuals in a patch, or equivalently when helping shows diminishing returns (Foster, 2004; Sibly and Curnow, 2011). In that case, being a rare type in a patch leads to fewer antagonistic interactions within patches. The selective advantage such a reduction brings depends on how often a type that is rare in the whole population is found in patches with individuals of a dissimilar type. This frequency is quantified in eq. (10) with *P*_*m*_, which is the probability that in a population where there is one common resident and one rare mutant type, two individuals randomly sampled in a mutant patch (a patch with at least one mutant) are not identical-by-descent (i.e. one is resident and the other is mutant, eq. C-21 in Appendix for details). Therefore when *P*_*m*_ is close to 0, a rare mutant tends to be found in patches showing low genetic variance (composed of mostly one type), whereas when *P*_*m*_ is larger, genetic variance within mutant patches is greater, leading to more interactions between mutant and resident within patches. *P*_*m*_ is greatest for intermediate dispersal rate *m* (Supp. Fig. S2) as when dispersal is extremely limited (*m* close to 0), then a mutant is mostly found in patches with other mutants, whereas when dispersal is unlimited (*m =* 1), a mutant is found only with residents.

The total effect of the dispersal rate *m* on mutual invasion, however, is not straightforward. In addition to influencing *P*_*m*_, limited dispersal on the one hand reduces the strength of selection by increasing kin competition (captured by (1−(1−*m*)^2^) in eq. 10), and on the other hand boosts selection under ecological inheritance as it increases the probability that future kin experience one’s environmental modifications (captured by Γ_*m*_ in eq. 10). Overall, we observe that −*I* is greatest and thus coexistence best protected when ecological inheritance *λ* is large and dispersal rate *m* is intermediate (Fig. 1C). In the absence of ecological inheritance *λ =* 0, however, −*I* always decreases with limited dispersal, highlighting the importance of ecological inheritance for the maintenance of variation in trait *z*.

### 3.2 Polymorphism drives long-term environmental heterogeneity due to ecological inheritance

When eq. (10) is negative, two weakly diverged types close to the evolutionary equilibrium *z*^*^ are maintained. If in addition selection is disruptive, these two types will diverge from one another, leading to a discrete polymorphism in a process known as “evolutionary branching” (Geritz et al., 1998). We show that this occurs in our model when the following holds,

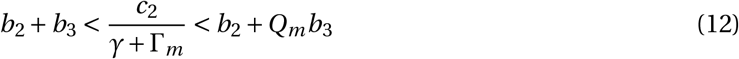

(Appendix C.3-C.6 for derivation). The inequality on the left hand side gives the condition for *z*^*^ to be convergence stable (eq. C-16 in Appendix C.3.3 for details) and the inequality on the right hand side gives the condition for *z*^*^ to be invadable (i.e. for selection to be disruptive; eq. C-18 in Appendix C.4 for details). Whether selection is disruptive depends on the compound parameter,

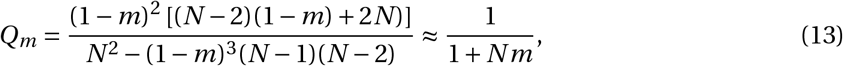

which reflects the genetic structure of the population. More specifically, *Q*_*m*_ is a measure of how common locally is a globally rare mutant: when *Q*_*m*_ *=* 1 (which occurs when *m =* 0), a rare mutant type finds itself only with other mutants; and conversely, when *Q*_*m*_ *=* 0 (which occurs when *m =* 1), it is only found with residents (eq. C-19 in Appendix for details). When *Q*_*m*_ *=* 1, the outer left and right hand side of inequality (12) are equal, so that evolutionary branching is impossible. This is to be expected since coexistence is impossible in that case (as *I* – eq. 10 – is equal to 0 when *P*_*m*_ *=* 0 owing to *m =* 1). By contrast, evolutionary branching is possible when *Q*_*m*_ *<* 1 (i.e. *m >* 0). In this case, evolutionary branching is facilitated by greater ecological inheritance (i.e. by greater Γ_*m*_) when *c*_2_ *>* 0 and *b*_2_ *+ b*_3_ *<* 0. These conditions correspond to a situation where the costs of investing into the local environment increase more than linearly with individual investment (i.e. accelerate, recall eq. 5) and the environmental benefits show diminishing returns with average investment in the patch (i.e. saturate, see eq. A-5 in Appendix).

We used individual-based simulations to investigate the outcome of evolutionary branching (Appendix C.7 for simulation procedure). These simulations show that the population becomes dimorphic with the gradual emergence and eventual maintenance of two extreme types: free-riders that express *z =* 0; and helpers that express *z = z*_max_ (recall eq. 3; Fig 2A). The emergence of this polymorphism is accompanied with an increase in variance among patches in average investment (within-patch average *z*), and consequently also in environmental quality *ϵ* (Fig. 2B). This heterogeneity is due to local sampling effects because of finite patch size, such that the average investment in a patch can fluctuate between 0 (only free-riders) and *z*_max_ (only helpers) over time (Supp. Fig. S3).

**Figure 2:**
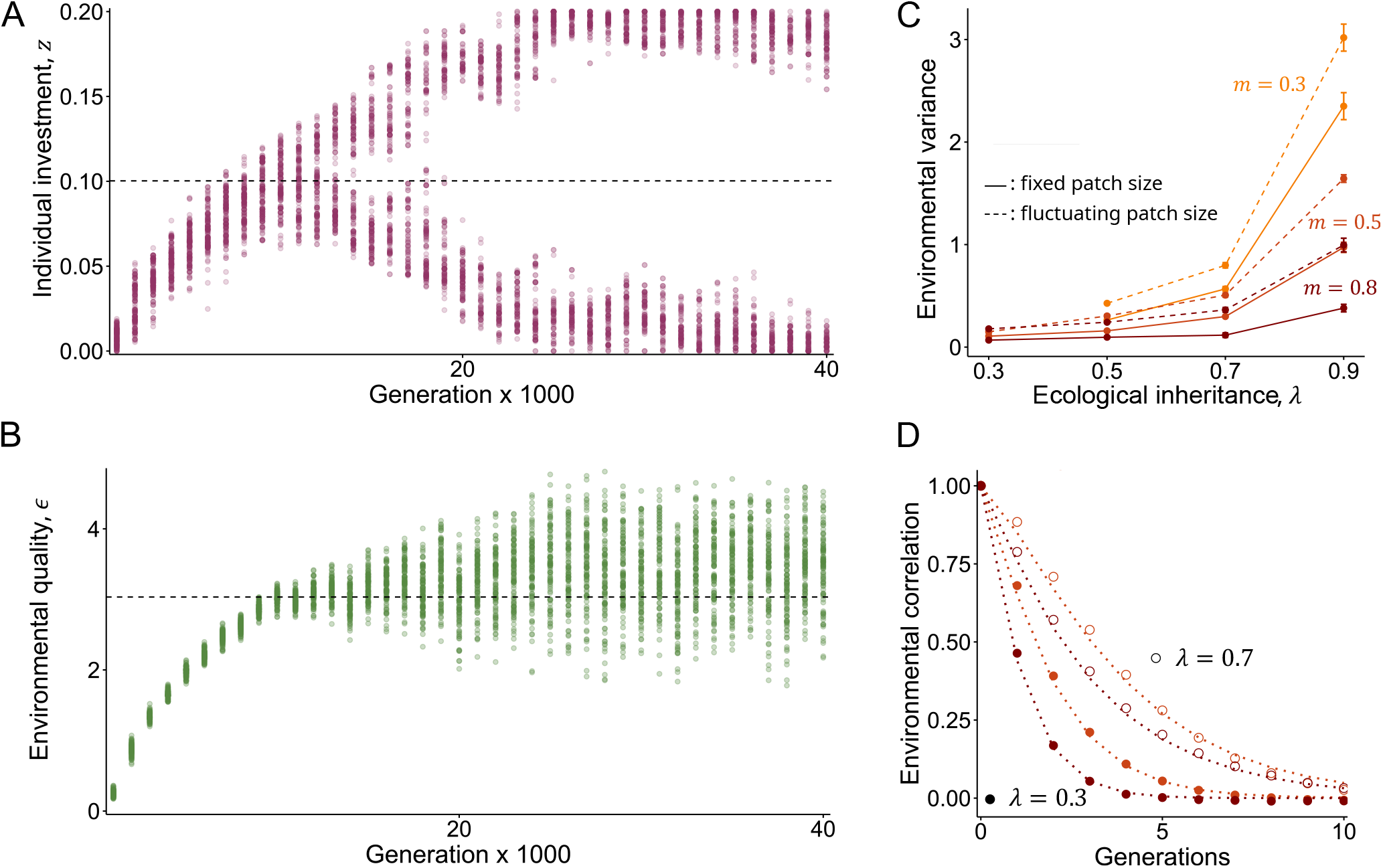
Evolutionary emergence of polymorphism and its ecological consequences. **(A-B) Evolution of helping and corresponding environmental dynamics**, from individual-based simulations (Appendix C.7 for procedure). (A) shows the trait values *z* of 150 randomly sampled individuals every 1’000 generations; while (B) shows the *ϵ* of 100 randomly sampled patches every 1’000 generations. These show the emergence of environmental helpers and free-riders (expressing *z* of approx. 0.2 and 0 respectively) and the resulting increase in environmental variance. The dotted line in (A) gives the evolutionary equilibrium *z*^*^ (eq. 8) and the dotted line in panel (B) gives the corresponding environmental variable 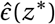 (eq. 9). **(C) Equilibrium polymorphic-driven environmental variance**. Among-patch variance in *ϵ*, averaged over 10 generations separated by 1’000 generations once polymorphism is established in simulations. Different colours are for different values of *m* (see legend). This shows that environmental heterogeneity is greater when *λ* is large and *m* is limited. Solid line: with fixed patch size (*N =* 10). Dashed line: with fluctuating patch size (such that when the population is monomorphic for *z*^*^, E[*N*] *=* 10; Appendix D for details). **(D) Equilibrium between-generation correlation in environment within patches**. Correlation Corr[*ϵ*_*t*_, *ϵ*_*t*−*h*_] within patches against *h* (x-axis), averaged over 50 generations once polymorphism is established in simulations. Different colours are for different values of *m* (see legend in C), full circles for *λ =* 0.3, empty circles for *λ =* 0.7. Dotted line: expected correlation, *ρ*_*ϵ,h*_, satisfying the recursion *ρ*_*ϵ,h+*1_ *= λρ*_*ϵ,h*_ *+* (1 − *λ*)(1 − *m*)^*h+*1^, with *ρ*_*ϵ*,0_ *=* 1. This shows that between-generation correlation is greatest when *λ* is large, following expectation. Other parameters: Panels ABCD: *b*_1_ *=* 1, *b*_2_ *=* 1.6, *b*_3_ *=* −2.5, *c*_2_ *=* 1.5, *γ =* 1. Panels AB: *λ =* 0.7, *m =* 0.5, *c*_1_ *=* 0.96. In panels CD, for each set of parameters the value of *c*_1_ is such that *z*^*^ *= z*_max_/2 *=* 0.1 across treatments to avoid boundary effects on variance and aid comparisons (specifically, *c*_1_ *=*(*γ+* Γ_*m*_) *b*_1_ − *z*_max_(*c*_2_ − (*γ+* Γ_*m*_) (*b*_2_ *+b*_3_))).

Environmental heterogeneity is especially high when ecological inheritance *λ* is large (Fig. 2C). This is because environmental effects are carried over multiple generations when *λ* is large and can therefore accumulate over time. This can be seen in Fig. 2D which shows the between-generations correlation in environmental quality within a patch (i.e. Corr[*ϵ*_*t*−*h*_, *ϵ*_*t*_] for different *h*) being greater when *λ* is large. Eventually, patches that have seen a succession of generations with many helpers are of much greater quality than in the absence of ecological inheritance; and conversely, patches with successive generations of free-riders remain of poor quality. Accordingly, limited dispersal (i.e. *m* small) enhances environmental heterogeneity as limited dispersal increases the between-generations correlation in the genetic composition of a patch (i.e. it increases intergenerational relatedness, Supp. Fig. S4, which entail successive generations of individuals in the same patch expressing similar traits).

### 3.3 Demographic fluctuations amplify environmental heterogeneity

The above results hold under the assumptions that all patches are of the same constant size *N*. However, among-patch variation in environmental quality could conceivably lead to among-patch demographic variation, and this would in turn feed back on the evolution of environmental investment. To investigate such feed back, we extended our simulations such that: (a) during stage (ii) of the life cycle (recall section 2.1), individuals now produce a random number of offspring whose distribution is Poisson with mean given by eq. (4); and (b) during stage (iv) of the life cycle each offspring is recruited as an adult with a density-dependent probability (Appendix D for details). To facilitate comparisons with results so far, we scale fecundity (eq. 4) so that the expected patch size in a population monomorphic for *z*^*^ is equal to patch size *N* in our simulations with fixed demography (i.e. as in Fig 2). As a preliminary check, we performed simulations using parameters such that condition (12) holds and such that it does not. We find that whether evolutionary branching occurs is well predicted from condition (12) (Supp. Fig. S5), suggesting that the emergence of polymorphism is not affected by demography here.

When polymorphism does emerge, it is accompanied with a change in population size albeit moderate (Fig. 3AB) that depends on whether within-patch trait variance increases or decreases patch quality. Specifically, when *b*_3_/(*N* − 1) *< b*_2_ then within-patch trait variance increases patch quality (eq. A-5 in Appendix A.2), which in turn augments fecundity and thus leads to a demographic boost (Fig. 3A). The left hand side of this inequality is always negative since *b*_3_ *<* 0 is necessary for evolutionary branching. When the right hand side, *b*_2_, is also negative, this indicates that the marginal effect an individual has on the environment saturates (recall eq. 2). As a result, the total effect of all patch members on the quality of their patch (ignoring interaction effects) is less than the sum of the marginal effects of each patch member. Within-patch trait variance thus tends to depreciate patch quality and thus fecundity. In contrast, if *b*_2_ *>* 0 then within-patch trait variance has a positive effect on fecundity (Fig. 3B).

**Figure 3:**
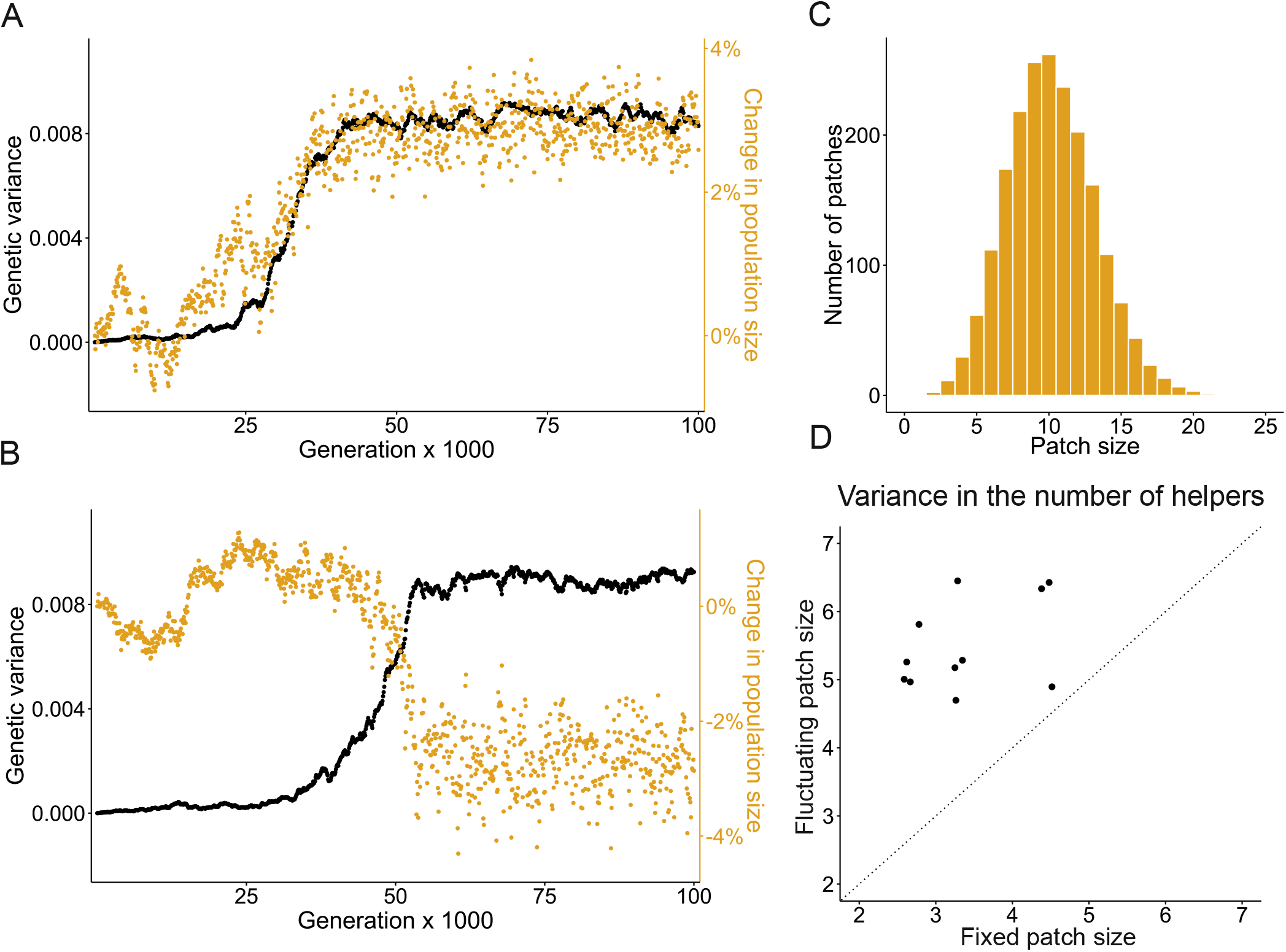
Demographic consequences of coexistence of environmental helpers and free-riders. **(A-B) Trait variance (black) and concomitant change in population size (yellow)** where the population size increases with polymorphism and ensuing trait variance (in A, with *b*_2_ *=* 5, *c*_2_ *=* 6.5) and where the population size decreases with polymorphism (in B, with *b*_2_ *=* −5, *c*_2_ *=* −10). From individual based simulation with fluctuating demography (Appendix D for details). Other parameters: *b*_3_ *=* −2.5, *b*_1_ *=* 1, *λ =* 0.7, *m =* 0.5, and *c*_1_ such that *z*^*^ *= z*_max_/2 *=* 0.1. **(C) Distribution of the number of adults per patch at equilibrium** (averaged over 20 observations separated by 1’000 generations). Parameter values: *b*_2_ *=* 1.6, *c*_2_ *=* 1.5. Other parameters: same as A. **(D) Variance in the number of helpers per patch under fixed vs. fluctuating demography**. This shows that the variance in helping per patch is larger when patch size fluctuates. Helpers are individuals with trait *z > z*^*^. Each data point is the variance observed in simulations where patch size is fixed (*N =* 10) vs where it fluctuates (with parameters such that E[*N*] *=* 10 when the population is monomorphic for *z*^*^) for a set of values for *m* and *λ* (Supplementary Table I for values used). Other parameters: same as C.

We observe appreciable variation in size among patches (Fig. 3C). To identify the drivers of this demographic heterogeneity, we calculated the correlations between different factors in the parental generation that can influence the number of offspring that will compete in that patch (namely: number of adults, environmental quality, and individual investment) and the size of the patch in the next generation (i.e. the number of offspring surviving after density-dependent regulation). These correlations are positive but typically weak (Supplementary Table I), likely because density-dependent regulation tends to smooth out demographic differences among patches. Accordingly, differences in patch size are mostly explained by local sampling effects, whereby chance effects determine whether more or less offspring survive to the next generation in each patch.

Polymorphic populations also show substantial variation in patch quality *ϵ* (dashed lines in Fig. 2C); in fact this variation is greater when patches vary in size than when size is fixed, especially when *λ* is large (compare solid lines and dashed lines in Fig. 2C). This indicates that demographic fluctuations amplify the effects of ecological inheritance on environmental heterogeneity. This is because the variance in the number of helpers per patch is greater when patches fluctuate in size compared to when patch size is fixed at *N* (Fig. 3D). With ecological inheritance, a patch that has been home to multiple generations of many helpers can then end up being of much greater quality than a patch home to multiple generations of free-riders.

### 3.4 Temporal and spatial heterogeneity are exacerbated by isolation-by-distance

Up until now, we have assumed that each patch is equally likely to be reached from any other patch by dispersal. As a final extension, we relax this assumption and incorporate isolation-by-distance into our model with patches now arranged on a lattice such that individuals can disperse only to neighbouring patches (“stepping-stone dispersal”, Kimura, 1953). Where patches are home to a single individual and individuals produce a public good that can diffuse across the whole lattice, isolation-by-distance can facilitate the coexistence of producers and non-producers (as spatial sorting can allow non-producers to exploit nearby producers; Wakano et al., 2009; Allen et al., 2013; Borenstein et al., 2013; Scheuring, 2014; Gerlee and Altrock, 2019). In our model, individuals contribute to the long term state of their patch (rather than neighbouring spots through diffusion); and patches can be home to an arbitrary number *N* of individuals (instead of just one). Previous analyses of such a scenario have shown that isolation by distance can boost positive selection on inter-generational effects (Lehmann, 2008; Mullon et al., 2024). In fact, using the results of these previous papers, the evolution of helping in our model under directional selection can be understood from eqs. (6) and (8) but with

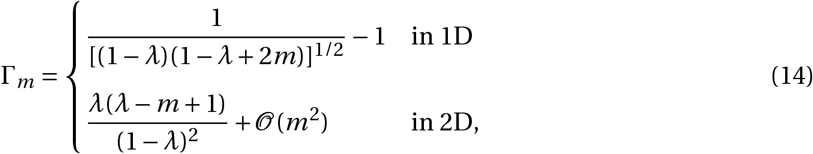

in a one- and two-dimensional lattices, respectively (Appendix E.1 for details). Both quantities in the right hand side of eq. (14) are greater than the right hand side of eq. (7), indicating that directional selection due to inter-generational effect is stronger under isolation-by-distance. This is because here, individuals that disperse can nevertheless land in patches that were previously inhabited by their ancestors and thus benefit from the contribution of these to the state of the patch. Intergenerational helping therefore evolves more readily under isolation-by-distance. But what of polymorphism?

We used individual-based simulations to investigate the establishment and consequences of polymorphism under isolation-by-distance, asking first whether isolation-by-distance facilitates the emergence of helpers and free-riders through evolutionary branching. To do so, we ran simulations for different values of *λ* (degree of ecological inheritance) and *m* (dispersal) under isolation-by-distance (on a two-dimensional 50*×*50 lattice) and under the island model. To avoid boundary effects on phenotypic variance, we adjusted the parameter *c*_1_ such that *z*^*^ remains the same across all simulations. Supplementary Figure S5 shows the phenotypic variance at mutation-selection-drift equilibrium, where low variance indicates a population that is essentially monomorphic whereas high variance indicates polymorphism. This reveals that the conditions for polymorphism are generally more restrictive under isolation-by-distance, requiring greater levels of dispersal, especially when ecological inheritance is weak (Fig. S5A). This is because competition among kin, which tends to inhibit polymorphism (Ajar, 2003; Wakano and Lehmann, 2014; Schmid et al., 2024), occurs more readily under isolation-by-distance (as such interactions can occur even among dispersers that land in the same patch) and therefore require greater levels of dispersal to be offset.

As before, a polymorphic population consists of essentially two types: free-riders that express *z =* 0; and helpers that express *z = z*_max_. To gain insights into the distribution of these types in time and space, we calculated the average genetic correlation between pairs of individuals that: (i) inhabit the same patch but at different generations (“within-patch” relatedness); and (ii) inhabit different patches but at the same generation (“between-patch” relatedness). As expected, within-patch relatedness decreases with generations (Fig. 4A) and between-patch relatedness with spatial distance (Fig. 4B). However, the decrease in within-patch relatedness with generation is slower when *λ* is large, i.e. individuals separated in time tend to be more related and thus more phenotypically similar when ecological inheritance is strong. This is because the environment tends to be more stable and less sensitive to genetic fluctuations in this case. As a result, selection on helpers and free-riders tends to be more homogeneous and thus more likely to maintain a stable proportion of each type within patches over time. This is reinforced by isolation-by-distance such that nearby patches show more similar genetic compositions – and far away patches more dissimilar compositions – when ecological inheritance is strong (as indicated by variation in between-patch relatedness being greater with spatial distance when *λ* is large, Fig. 4B).

**Figure 4:**
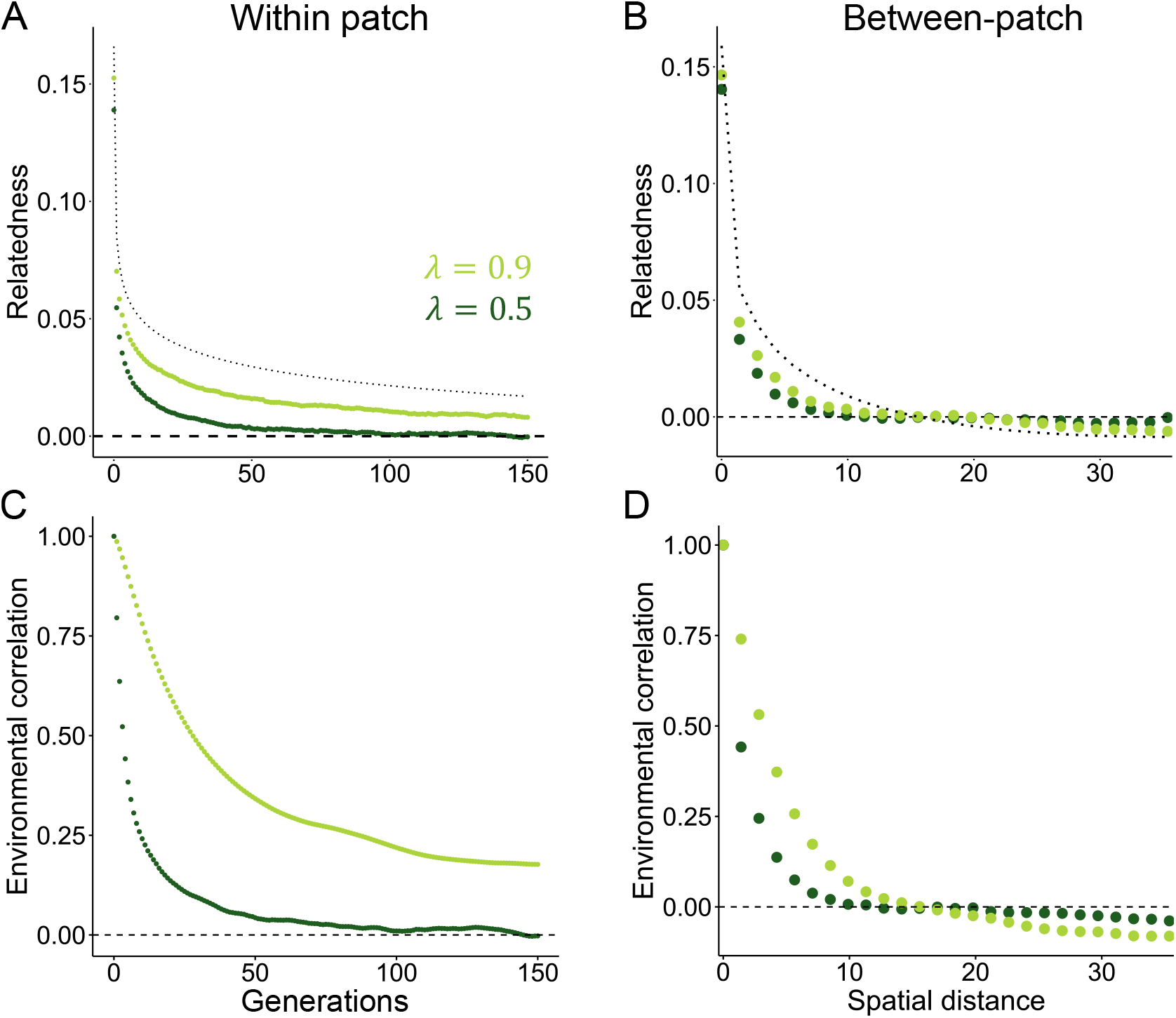
Genetic and environmental structure in polymorphic populations under isolation-by-distance. **(A) Inter-generational relatedness**, i.e. average correlation in mean trait value in a patch between generations. **(B) Inter-patch relatedness**, i.e. average correlation in mean trait value between patches at the same generation. Dotted lines in A-B: relatedness in a monomorphic population (from eq. 8 in Mullon et al., 2024). **(C) Average correlation in the environment of a patch between generations. (D) Average correlation between the environments of different patches at the same generation**. Dark green: *λ =* 0.5. Light green: *λ =* 0.9. Other parameters: *b*_1_ *=* 1, *b*_2_ *=* 2, *b*_3_ *=* −2.5, *c*_2_ *=* 1.5, *m =* 0.8, *N =* 10, *c*_1_ chosen such that *z*^*^ *= z*_max_/2 *=* 0.1. Appendix E.2 for simulation details.

The spatio-temporal patterns of helpers and free-riders are reflected in the distribution of environmental states in time and space. Specifically, the correlation between the environmental states of the same patch at different generations is greater (Fig. 4C), while between-patch correlation varies more substantially with spatial distance (Fig. 4D), when ecological inheritance is strong. In addition, environmental correlations are consistently greater than genetic correlations under ecological inheritance (compare Figs. 4AB with Figs. 4DE). Overall, these results show that under the combined effects of isolation-by-distance and ecological inheritance, genetic and environmental fluctuations interact. This interaction leads to more structured environmental heterogeneity in space and time, with the two-dimensional landscape showing clusters of “good” and “bad” patches that change more slowly when *λ* is large (Fig. 5 and supplementary movies available in the online supplementary material, Prigent and Mullon, 2025).

**Figure 5:**
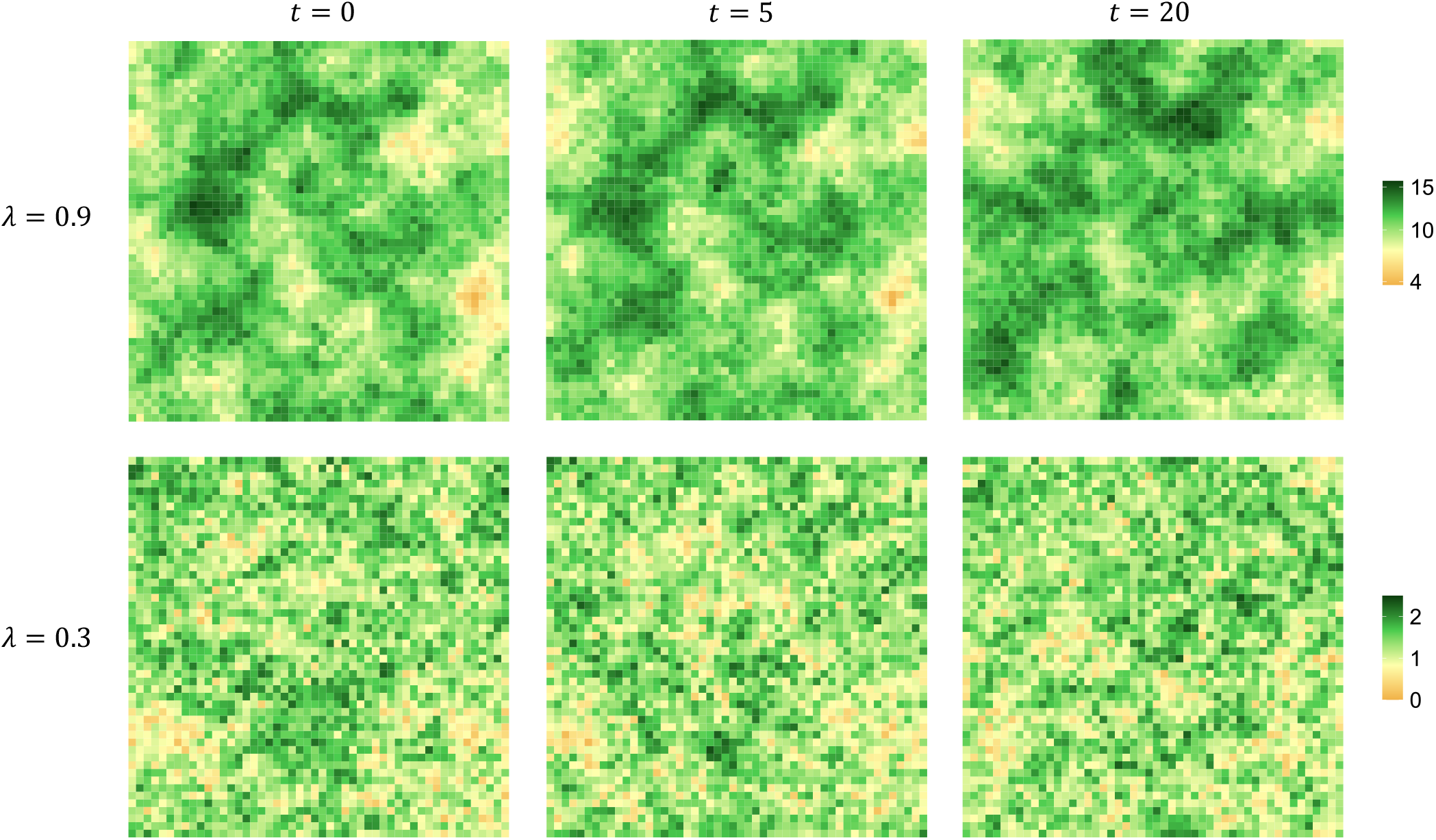
Polymorphism-driven spatio-temporal dynamics of the environment. Each panel shows the environmental variables of all patches of a polymorphic population at different time points. See figure legend for scale. Top panel: *λ =* 0.9. Bottom panel: *λ =* 0.3. Other parameters: same as Fig. 4. Appendix E.2 for simulation details.

## 4 Discussion

Our analyses indicate that ecological inheritance, when coupled with limited dispersal, promotes the emergence and maintenance of polymorphism in environmentally mediated helping behaviours. This is because these conditions increase the growth rate of both helpers and free-riders when they are rare. Under the combination of ecological inheritance and limited dispersal, the environmental improvements made by a rare lineage of helpers persist across generations and preferentially benefit their philopatric descendants; a rare lineage of free-riders meanwhile can temporarily reap the rewards of others’ contributions without incurring the costs of defection (at least initially). Our findings contrast with previous results in well-mixed populations, where ecological inheritance alone has been shown to destabilise polymorphisms and drive cyclical or runaway dynamics (Brown and Taddei, 2007; Weitz et al., 2016; Tilman et al., 2020, see Appendix F for further discussion). Given that many natural populations are dispersal-limited, our results suggest that ecological inheritance may help maintain variation in socially relevant traits such as nest construction, burrow excavation, and resource accumulation—behaviours that have been reported to exhibit marked inter-individual variation (Clutton-Brock et al., 2003; Thorley et al., 2018; Siegmann et al., 2021; Jeanson and Weidenmüller, 2014). However, the extent to which these environmental effects influence the fitness of future generations remains to be characterised.

Animal societies provide a useful context to test these ideas, as they often exhibit substantial relatedness. In social insects, genetic variation within colonies has been linked to helping behaviours that modify the environment. In wasps, genetic markers correlate with differences in nest-building effort (O’Donnell, 1996). In honeybees, genotypic differences influence wax production and comb construction (Calderone and Page, 1988; Robinson, 1989; Page and Robinson, 1991; Dreller et al., 1995), while in ants, genetic factors contribute to variation in nest maintenance and tunnel excavation (Julian and Fewell, 2004). In social mammals, the available data suggest that environmentally mediated helping is more strongly influenced by individual condition than by genetics (e.g. Clutton-Brock et al., 2003; Thorley et al., 2018; Siegmann et al., 2021 for habitat maintenance). Nevertheless, social traits in mammals typically exhibit significant additive genetic variance (Houslay et al., 2021; Nichols et al., 2021), suggesting that selection could readily drive the evolution of environmentally mediated helping behaviours.

Microbial and plant systems may also provide productive empirical avenues to test how ecological inheritance influences the persistence of helping traits. In *Pseudomonas aeruginosa*, experimental manipulations of the durability of pyoverdin show that more persistent forms allow cooperators to better withstand exploitation by free-riders (Kümmerli and Brown, 2010), supporting the idea that long-lasting environmental modifications can stabilise helping behaviour. Meanwhile, many plant species produce root exudates and litter that enrich the soil, and because seeds often disperse only short distances, offspring often inherit these improvements alongside their associated microbial communities (Bever, 1994, 2002; Van der Putten et al., 2016). If there is heritable variation in these soil-modifying traits, certain plant lineages may gain lasting benefits from enhanced rhizospheres, whereas genotypes that invest less could exploit soil enriched by others. For instance, empirical studies reveal genetic differences in exudate composition and litter quality that significantly affect microbial activity and plant performance (e.g. Badri and Vivanco, 2009). Further research tracking how such differences influence soil conditions and plant fitness across generations could clarify the role of ecological inheritance in maintaining trait variation.

In humans, long-term environmental decision-making varies widely between individuals, influencing behaviours such as resource conservation, land use, and responses to large-scale environmental challenges like climate change (Milfont and Gouveia, 2006; Carmi, 2013). While this variation is strongly influenced by socio-economic background and education (Rickinson, 2001; Bayard and Jolly, 2007), there is also evidence for heritable differences in personality traits (Bouchard, 2004; Kandler, 2012). Many of these traits indirectly affect environmental concern and pro-environmental behaviour; for example, individuals who are more open to experience tend to be more receptive to new ideas and change, which may foster stronger environmental attitudes and actions (Hirsh, 2010). Our model highlights one way in which such genetic variation may be maintained by selection.

Variation in environment-modifying traits has significant ecological—and therefore fitness—consequences. Two key processes shape these effects in our model. First, once polymorphism is established, local sampling effects under limited dispersal generate environmental heterogeneity among patches (see Bolnick et al., 2011 for a general discussion on the ecological consequences of sampling effects). This effect is amplified by ecological inheritance, as the environment reflects not only the contributions of the current generation but also the cumulative effects of past generations. Second, the relationship between individual trait expression and environmental quality is often non-linear and non-additive, as parametrized by *b*_2_ and *b*_3_ (eq. 4; see also eq. A-5 in Appendix). By Jensen’s inequality (Ruel and Ayres, 1999; Bolnick et al., 2011), if *b*_2_ *>* 0 (such that the marginal effect of an individual’s trait expression on the environment accelerates with greater trait values) then trait variance within patches tends to improve local environmental conditions (whereas if *b*_2_ *<* 0, greater trait variance tends to degrade the environment). Stable polymorphism in our model requires negative interaction effects *b*_3_ *<* 0, meaning that individuals with different trait values interfere less with one another. This further strengthens the positive environmental effects of trait variation within patches, leading to differences in average environmental quality across subpopulations.

The assumption that individuals with different trait values interfere less with one another (*b*_3_ *<* 0), which is necessary for coexistence in our model, would naturally arise when helping has diminishing returns on environmental quality. Such saturating effects are thought to be widespread (see Foster, 2004, for a review). For instance, in the paper wasp *Polistes dominula*, increasing the number of helpers leads to diminishing benefits for nest growth (Grinsted and Field, 2018).

While interference provides one route through which long-term environmental modifications favours trait variation, it is not the only possible mechanism. In our supplementary material (Appendix G), we show that ecological inheritance mediates such selection effects through multiple pathways. One such pathway which may be particularly relevant in natural conditions occurs when the fitness benefits of environmental quality plateau beyond a certain threshold, as other factors become limiting (eq. G-6 in Appendix for details). For example, in the yeast *Saccharomyces cerevisiae*, individuals secrete invertase, an enzyme that hydrolyses sucrose into glucose and fructose, thereby increasing the availability of glucose in the environment. However, at high glucose concentrations, the efficiency of glucose uptake by cells declines due to a “rate-efficiency” trade-off (MacLean et al., 2010); so that beyond a certain level of invertase production, additional enzyme results in diminishing returns. Beyond this specific example, we find that via all pathways through which the environment mediates selection, variation in environment-modifying behaviours is favoured by the combination of ecological inheritance and limited dispersal (Appendix G for details). The prevalence of restricted gene flow suggests that these processes play a role in shaping long-term ecological and evolutionary dynamics, potentially driving the emergence of spatially structured ecosystems and sustained behavioural diversity in systems ranging from microbial biofilms and plant–soil interactions to social insect colonies and burrowing mammals.

## Supplementary Figures and Tables

**Supplementary Figure S1:**
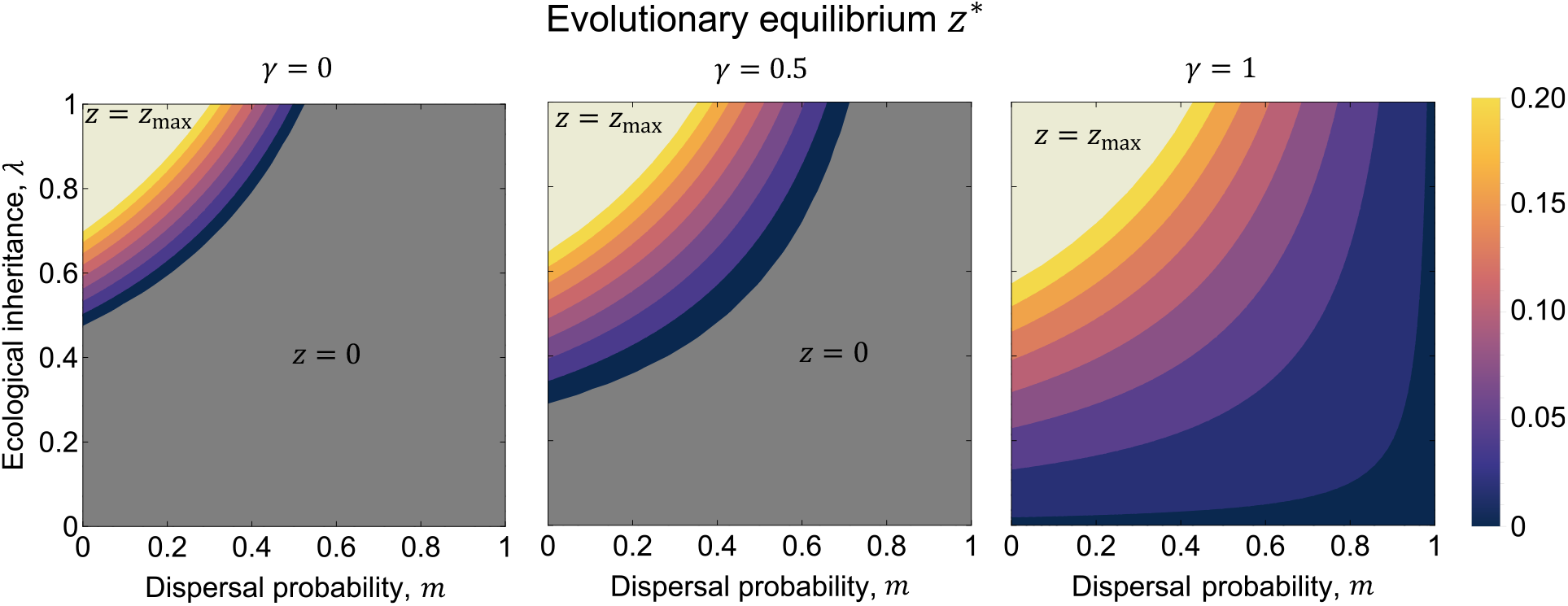
Evolutionary equilibrium of the investment into the environment, *z* ^*^. (from eq. 8). This shows that selection favours large *z*^*^ when dispersal *m* is low, ecological inheritance *λ* is high, and the proportion of individual investment that individuals get to enjoy in their own lifetime *γ* is high. The dark grey region shows the space of parameter for which investment into the environment does not evolve (i.e. inequality 6 is not satisfied). Other parameters: *b*_1_ *=* 1, *b*_2_ *=* 1.6, *b*_3_ *=* −2.5, *c*_1_ *=* 0.9, *c*_2_ *=* 1.5.

**Supplementary Figure S2:**
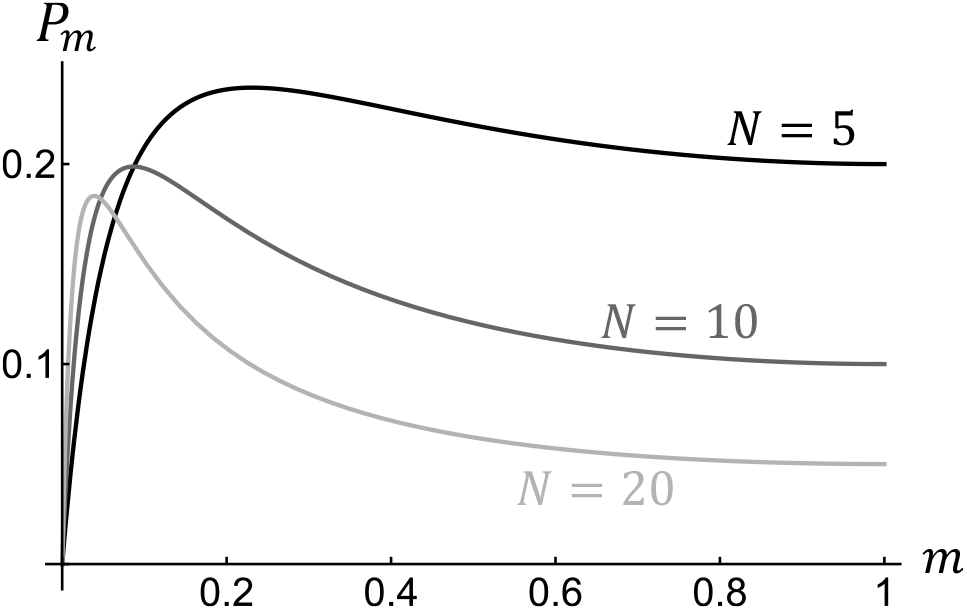
Probability *P*_*m*_ that two individuals sampled without replacement from a mutant patch are not identical-by-descent. (i.e., one is a mutant and one is a resident; eq. C-21). This probability is greatest for intermediate values of *m* where both mutants and residents are equally present in the patch.

**Supplementary Figure S3:**
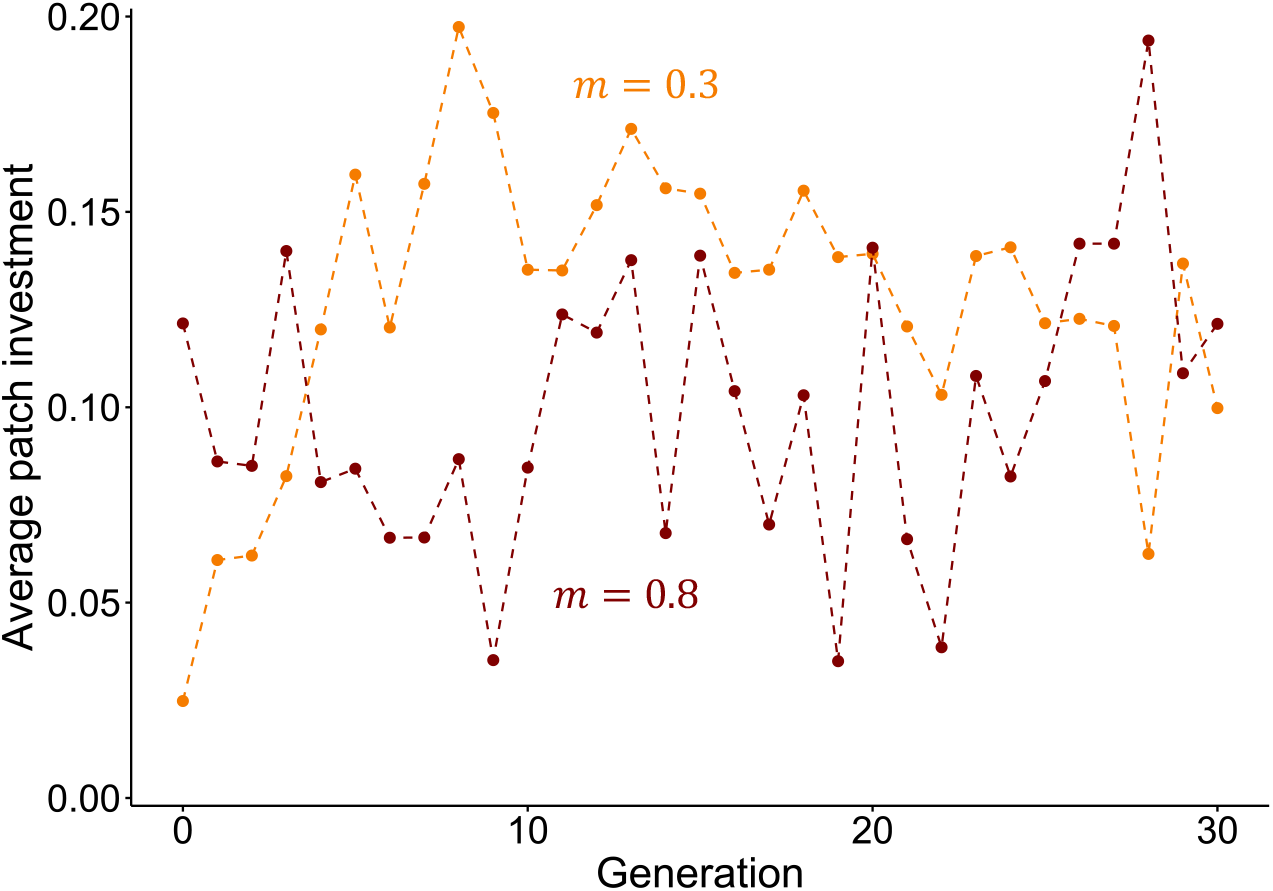
Average trait expressed in randomly chosen patch across generations in a polymorphic population. As each of the *N =* 10 individuals within a patch expresses either *z ≈* 0 or *z ≈ z*_max_ *=* 0.2, the average trait fluctuates between these two values. When dispersal *m* is greater, there is less correlation in average trait from one generation to the next 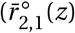 is lower, as shown is Fig. S4), meaning that the fluctuations in average trait across generation are greater. Different colours are for different values of *m* (see legend). Other parameters: *b*_1_ *=* 1, *b*_2_ *=* 1.6, *b*_3_ *=* −2.5, *c*_2_ *=* 1.5, *λ =* 0.7, *c*_1_ chosen such that *z*^*^ *= z*_max_/2 *=* 0.1. Appendix C.7 for simulation procedure.

**Supplementary Figure S4:**
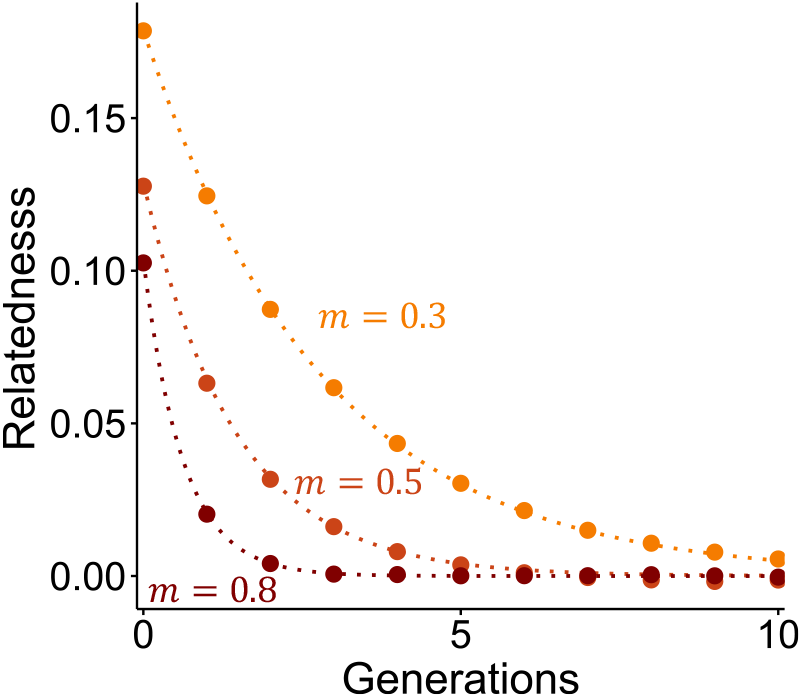
Inter-generational relatedness, 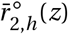, as a function of *h*. Dotted line: expected value of inter-generational relatedness under neutrality (eq. II.1). Dots: estimates from individual based simulations (computed as the correlation in the average trait value within patches separated by *h* generations – averaged over 50 generations once polymorphism is established in population). Different colours are for different values of *m* (see legend). Other parameters: *b*_1_ *=* 1, *b*_2_ *=* 1.6, *b*_3_ *=* −2.5, *c*_2_ *=* 1.5, *λ =* 0.7, *c*_1_ chosen such that *z*^*^ *= z*_max_/2 *=* 0.1. Appendix C.7 for simulation procedure.

**Supplementary Figure S5:**
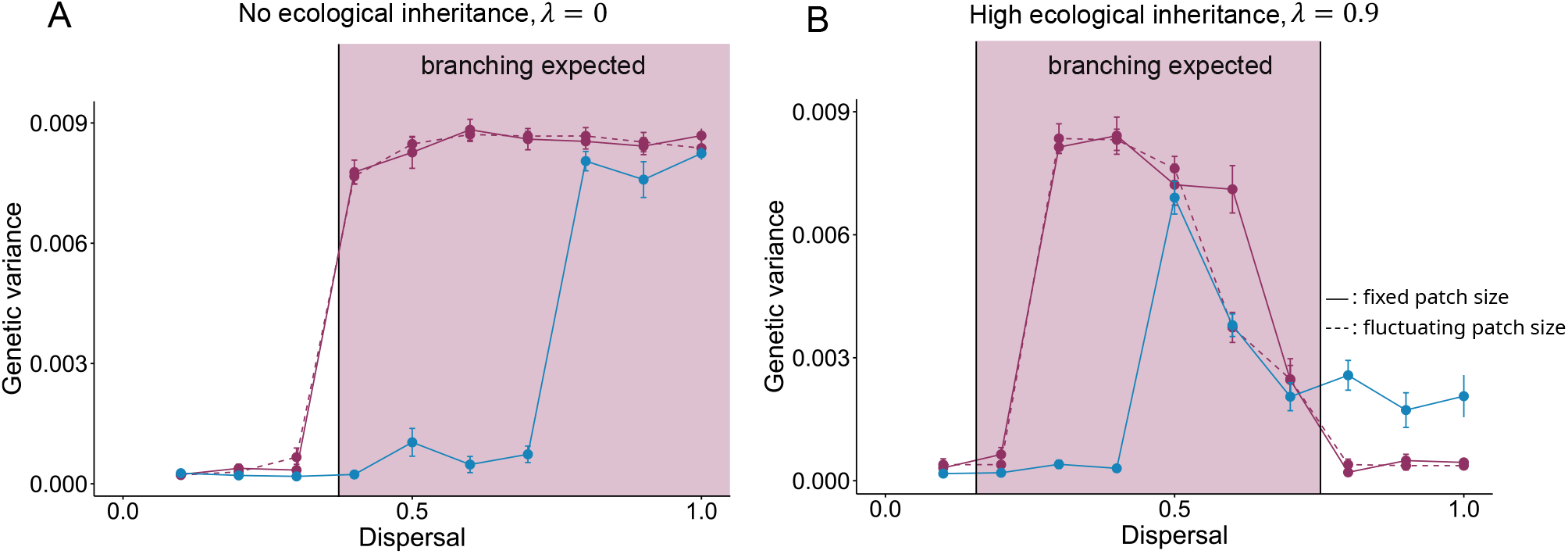
Equilibrium trait variance in the population, from individual-based simulations. Appendices C.7, D and E.2 for simulation procedures. The variance Var[*z*] is average over 10^*′*^000 generations (from generation 290^*′*^000 onwards, such that polymorphism has had time to establish if it is favoured by selection). In purple: the population evolves under the island model of dispersal (solid line, with fixed demography with patch size *N =* 10; dashed line, under fluctuating demography with average patch size E[*N*] *=* 10). Conditions for evolutionary branching are fulfilled under the island model of dispersal with fixed demography in the light purple region (i.e. where ineq. 12 holds). In blue: the population evolves under the stepping-stone model of dispersal in a 2D lattice. This shows that letting demography fluctuate has no discernible effect on the genetic variance (i.e., ineq. 12 predicts accurately whether evolutionary branching occurs or not), but that the conditions for evolutionary branching to occur are generally more restricted under isolation-by-distance. Parameters values: Panel A: *b*_2_ *=* 1.8, *λ =* 0.0. Panel B: *b*_2_ *=* 1.2, *λ =* 0.9. Other parameters: *b*_1_ *=* 1, *b*_3_ *=* −2.5, *c*_1_ *=* 0.9, *c*_2_ *=* 1.5, *c*_1_ chosen such that *z*^*^ *= z*_max_/2 *=* 0.1. Total number of patches: 2’500.

**Supplementary Figure S6:**
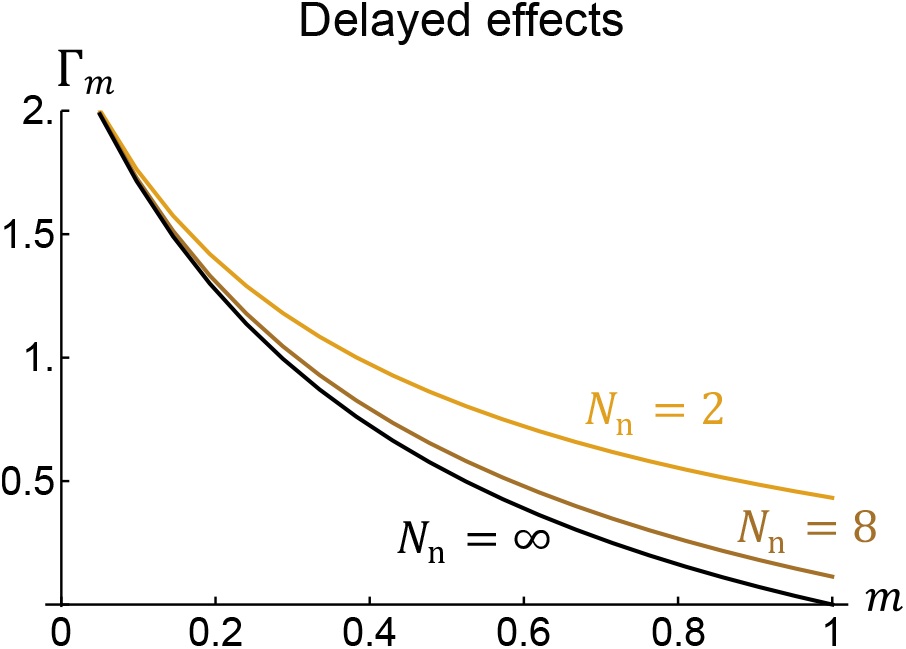
Strength of selection on inter-generational helping, Γ_*m*_, under isolation by distance. Results are shown for 1D lattice (with two neighbouring patches *N*_n_ *=* 2, from eq. E-17), 2D lattice (with *N*_n_ *=* 8, from eq. E-18), and for the island model (*N*_n_ *=* ∞ using eq. 7.

**Supplementary Table I:** Correlation between parental generation (patch size, patch quality, and trait values) and patch size in the following generation, from individual based simulations (Appendix D). This shows that factors in the parental generation only explain a small amount of the variation in patch size in the offspring generation, especially when dispersal *m* is high. Parameters values: *b*_1_ *=* 1, *b*_2_ *=* 1.6, *b*_3_ *=* −2.5, *c*_2_ *=* 1.5, E[*N*] *=* 10, *c*_1_ chosen such that *z*^*^ *= z*_max_/2 *=* 0.1. §: Evolutionary branching does not occur under this set of parameters.

| Dispersal | Ecological inheritance | Correlation between offspring patch size $N_{k,t+1}$ and parental | | |
| --- | --- | --- | --- | --- |
| | | patch size, $N_{k,t}$ | patch quality $\epsilon_{k,t}$ | trait values $z_{i,k,t}$ |
| $m = 0.3$ | $\lambda = 0.3^{\S}$ | $0.091 \pm 0.025$ | $0.031 \pm 0.028$ | $0.009 \pm 0.010$ |
| | $\lambda = 0.5$ | $0.078 \pm 0.028$ | $0.039 \pm 0.015$ | $0.009 \pm 0.008$ |
| | $\lambda = 0.7$ | $0.078 \pm 0.019$ | $0.029 \pm 0.020$ | $0.010 \pm 0.008$ |
| | $\lambda = 0.9$ | $0.072 \pm 0.022$ | $0.033 \pm 0.023$ | $0.004 \pm 0.010$ |
| $m = 0.5$ | $\lambda = 0.3$ | $0.064 \pm 0.021$ | $0.012 \pm 0.020$ | $0.008 \pm 0.008$ |
| | $\lambda = 0.5$ | $0.070 \pm 0.027$ | $0.022 \pm 0.016$ | $0.006 \pm 0.010$ |
| | $\lambda = 0.7$ | $0.061 \pm 0.018$ | $0.027 \pm 0.025$ | $0.005 \pm 0.007$ |
| | $\lambda = 0.9$ | $0.021 \pm 0.023$ | $0.021 \pm 0.023$ | $0.006 \pm 0.007$ |
| $m = 0.8$ | $\lambda = 0.3$ | $0.031 \pm 0.017$ | $0.002 \pm 0.030$ | $0.002 \pm 0.005$ |
| | $\lambda = 0.5$ | $0.016 \pm 0.024$ | $0.008 \pm 0.022$ | $0.003 \pm 0.006$ |
| | $\lambda = 0.7$ | $0.027 \pm 0.022$ | $0.001 \pm 0.031$ | $0.003 \pm 0.009$ |
| | $\lambda = 0.9$ | $0.010 \pm 0.025$ | $0.004 \pm 0.021$ | $0.001 \pm 0.006$ |

## Appendix

### A Some further details on the model

In this Appendix, we specify the effect of trait *z* on the environment in the model, complementing section 2.1 of the main text.

#### A.1 Investment is a form of helping

First, we show that eq. (3) of the main text must hold for *z* to always constitute a form of helping. For this to be the case, we need that *B* (***z***_*t*_) is always increasing with individual investment, i.e. that

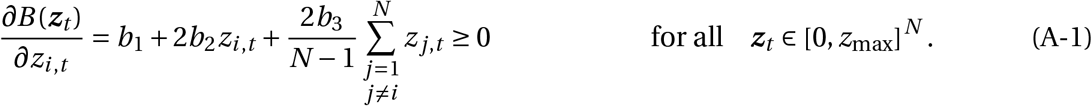

Since this must be true when *z*_*i,t*_ *=* 0 for all *i*, we must have *b*_1_ *>* 0. We then seek to characterise *z*_max_ such that eq. (A-1) always holds, which depends on *b*_2_ and *b*_3_.

When *b*_2_ *>* 0 and *b*_3_ *>* 0, eq. (A-1) holds for any 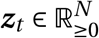 so that *z*_max_ → ∞. When *b*_2_ *>* 0 and *b*_3_ *<* 0, *∂B* (***z***_*t*_)/*∂z*_*i,t*_ increases with *z*_*i,t*_ but decreases with *z* _*j,t*_. Therefore *∂B* (***z***_*t*_)/*∂z*_*i,t*_ is at its minimum when *z*_*i,t*_ *=* 0 and *z* _*j,t*_ *= z*_max_ for all *j*≠ *i*, where it must be non-negative. Therefore we require that

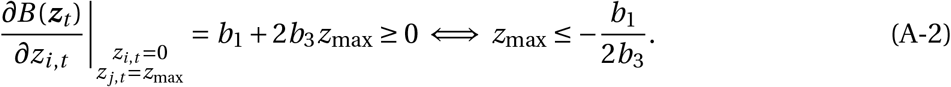

Conversely, when *b*_2_ *<* 0 and *b*_3_ *>* 0, *∂B* (***z***_*t*_)/*∂z*_*i,t*_ is at its minimum when *z*_*i,t*_ *=* 0 and *z* _*j,t*_ *= z*_max_, which is non-negative if and only if

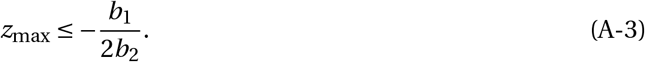

Finally, when *b*_2_ *<* 0 and *b*_3_ *<* 0, *∂B* (***z***_*t*_)/*∂z*_*i,t*_ is at its minimum when *z*_*i,t*_ *= z*_max_ for all *i*, and is non-negative if and only if

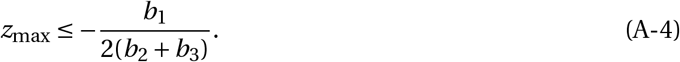

The above conditions give us eq. (3) of the main text, as required.

#### A.2 The effects of mean and variance in helping within groups

Eq. (2) expresses *B* (***z***_*t*_) in terms of the investment *z*_*i,t*_ of each individual *i* in a patch. It may be useful for interpretation purposes to express *B* (***z***_*t*_) in terms of the patch-mean 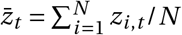 and variance 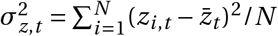 in investment, i.e. as

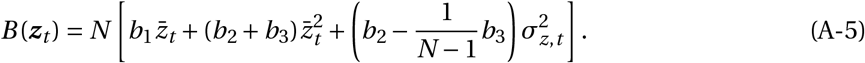

Eq. (A-5) makes it apparent that the production common good increases with trait variance in the patch if, and only if, *b*_2_ −*b*_3_/(*N* − 1) *>* 0 or equivalently *b*_3_/(*N* − 1) *< b*_2_.

### B Connections with models evolutionary game theory and niche construction

In this appendix, we detail how our model connects to evolutionary game theory models (in particular those with long term environmental effects, Weitz et al., 2016; Brown and Taddei, 2007; Tilman et al., 2020) as well as niche construction models (Laland et al., 1999; Silver and Di Paolo, 2006; Odling-Smee et al., 2003). But first note that when *λ =* 0 and *γ =* 1, all the effects of helping occur within generations, such that with *N =* 2, our model of social interactions reduces to standard 2-players continuous strategy games. The marginal payoff for an individual cooperating more is *∂B* (*z*_*i,t*_, *z* _*j,t*_)/*∂z*_*i,t*_ − *∂C* (*z*_*i,t*_)/*∂z*_*i,t*_, while the marginal payoff for the partner of an individual cooperating more is *∂B* (*z*_*i,t*_, *z* _*j,t*_)/*∂z*_*j,t*_. For instance, we recover a continuous Prisoner’s Dilemma with *b*_2_ *= c*_2_ *= b*_3_ *=* 0 and 0 *< b*_1_ *< c*_1_, and a Hawk-Dove or Snowdrift Game with 0 *< c*_1_ *< b*_1_ and *b*_3_ *<* −*b*_1_ (Rapoport and Chammah, 1965; Maynard Smith and Price, 1973; Maynard Smith, 1982). Our model can also recover *N* -player continuous games when *N >* 2. In particular, setting *b*_3_/(*N* − 1) *= b*_2_ recovers the model of Doebeli et al. (2004) (their SOM 1.1.3, where the gains are divided between *N* randomly chosen individuals, i.e. with *m =* 1; see also Wakano and Lehmann, 2014, their eq. 33-34 for this model under limited dispersal *m <* 1 under pairwise interactions).

In the presence of ecological inheritance (*λ >* 0), our eq. (1) can be seen as the discrete time equation of the ecological dynamics found in Brown and Taddei (2007) (their eq. 1) and in Tilman et al. (2020) (their eq. 9, with *e*_L_ *=* 0). In Weitz et al. (2016), ecological dynamics (their eq. 17) differ from those in our model only in that they assume that the environment is degraded only when free-riders are present (and not in their absence). Our eq. (1) also allows us to connect to niche construction models: eq. (1) in Laland et al. (1999), eq. (1) in Silver and Di Paolo (2006), eq. (A3.1) in Odling-Smee et al. (2003). All these are equal to our eq. (1) when they assume positive niche construction and no independent renewal of the niche (in their notation: 0 *< λ*_2_ *<* 1, *γ =* 0 and *λ*_3_ *=* 0).

We differ however from all these models on one relevant assumption about environmental effect on fitness. Whereas we assume that improvement in environmental quality has the same positive effect on an individual’s fecundity, regardless of its trait *z* (eq. 4), all these models assume that the environment affects fecundity via trait-by-environment effects. Specifically :

1. In niche construction models, it is assumed that niche construction is favourable only for those individuals carrying the appropriate recipient allele (otherwise it is harmful for those carrying the other allele, Table 1 Laland et al., 1996, eq. 2 in Silver and Di Paolo, 2006). Where the environmental effect of niche construction is global (i.e. common to the whole population), this entails that the invasion of an allele encoding for niche construction does not depend on its own environmental effect (Laland et al., 1996). Where the environmental effect of niche construction is local, invasion of the niche construction allele relies on its association to the appropriate recipient allele (Fig. 2 in Silver and Di Paolo, 2006). Importantly, polymorphism at the niche construction locus cannot be maintained in either case, unless this allele shows built-in overdominance (as assumed in Laland et al., 1996).
2. In evolutionary game theory models, meanwhile, it is assumed that free-riders have an advantage in good quality environments and helpers in bad quality environments (eq. 12 in Weitz et al., 2016, eq. 5 in Tilman et al., 2020, first matrix in the section ‘Durable Public good’ in Brown and Taddei, 2007). Biologically, this assumes that helpers are able to partially privatize their investment before it affects the shared environment. This assumption is necessary for helping to evolve where environmental effects are global (i.e. common to the whole population). Otherwise, free-riders (or ‘cheaters’) always have a greater fitness that helpers, leading to a form of the tragedy of the commons; as described e.g. in eq 1-7 in Weitz et al., 2016).

We discuss further the implications that these differences in assumptions have in a supplementary discussion (Appendix F).

### C Evolutionary dynamics under the island model of dispersal with constant demography

In this appendix, we investigate our model under the baseline assumptions that dispersal is uniform among patches and that each patch is of fixed size *N*. In particular, we derive eqs. (6)-(13) of the main text.

#### C.1 General approach

We are interested in whether ecological inheritance (modulated by the parameter *λ*) either promotes or inhibits polymorphism in trait *z*. Assuming that *z* evolves via the input of rare genetic mutations with weak phenotypic effects, evolutionary dynamics can be determined via an analysis of the invasion fitness *W* (*ζ, z*) of a rare mutant with trait value *ζ* arising in a resident population that is monomorphic for trait values *z* (Geritz et al., 1998; Rousset, 2004; Dercole and Rinaldi, 2008; Avila and Mullon, 2023).

Trait evolution unfolds in two tempos under these assumptions. First, the trait evolves under directional selection whereby positively selected mutants rapidly sweep to fixation such that the population effectively jumps from one monomorphic state to another. The direction of selection is determined by the selection gradient

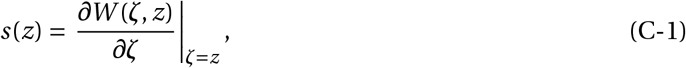

with the trait increasing when *s*(*z*) *>* 0 and decreasing when *s*(*z*) *<* 0. The population may eventually reach an evolutionary equilibrium *z*^*^ such that

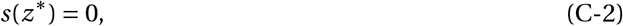

and

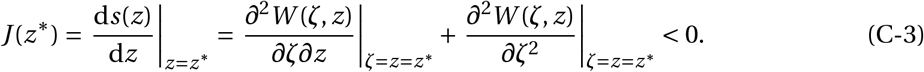

Such a *z*^*^ is referred to as convergence stable. Second, the population either remains monomorphic for a convergence stable *z*^*^ under stabilising selection, or becomes polymorphic due to disruptive selection in a process referred to as evolutionary branching (Geritz et al., 1998). This is determined by the sign of

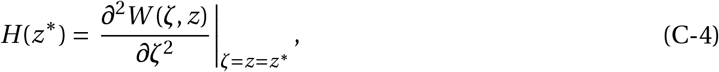

with *H* (*z*^*^) *<* 0 indicating stabilising selection and *H* (*z*^*^) *>* 0 indicating disruptive selection.

From the above, the emergence of polymorphism requires that the following condition holds,

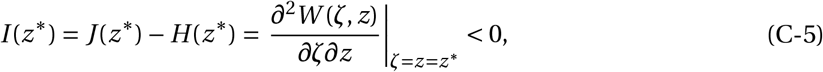

i.e., that the invasion fitness of a mutant decreases with joint changes in the trait of its own lineage and in the resident trait. When condition (C-5) holds, invasion fitness increases as a result of mutant and resident trait values diverging away from *z*^*^. The cross derivative *I* (*z*^*^) can therefore be seen as trait-dependent selection, and such selection must be negative for polymorphism (Lehmann and Mullon, 2025). In fact, two types coding for *z*^*^ *+* Δ and *z*^*^ − Δ can reciprocally invade one another (see eq. A1 in Geritz et al., 1998 or eq. A1 in Lehmann and Mullon, 2025) – a scenario consistent with negative frequency-dependent selection – if and only if *I* (*z*^*^) *<* 0 when Δ is small.

For the island model of dispersal, previous studies have derived the selection gradient *s*(*z*) (Lehmann, 2007) and the disruptive selection coefficient *H* (*z*) (Prigent and Mullon, 2023) on traits that have long-term local environmental effects (see also Ohtsuki et al., 2020 for a general expression for *s*(*z*) and *H* (*z*) in non-homogeneous subdivided populations, where the non-homogeneity can be driven by environmental effects of the evolving trait). We use these results to derive *s*(*z*) and *H* (*z*) for our model, and in turn use eq. (C-5) to investigate trait dependent selection *I* (*z*).

#### C.2 Building blocks

First, we specify the necessary building blocks to obtain *s*(*z*) and *H* (*z*), using the notation of Prigent and Mullon (2023) (their sections 2-3) to facilitate applying their results.

##### C.2.1 Environmental effects

The first building block is the environmental map *F*, which gives the environmental transformation from one generation to the next. According to our assumptions, this is,

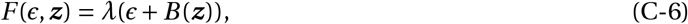

where *B* is given by eq. (2) of the main text (recall eq. 1 in main text). In the absence of trait variation (when all individuals express *z* so that ***z*** *=* (*z*,…, *z*)), all patches converge to the same environmental equilibrium, which is such that 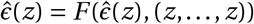, i.e.

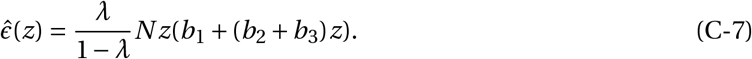

provided

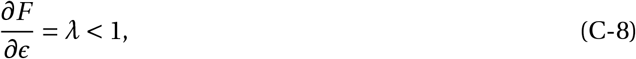

which is assumed throughout. To apply the results of Prigent and Mullon (2023), it will be useful to also define

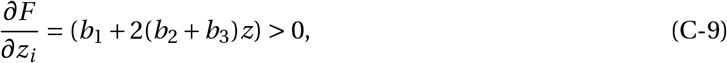

where the derivative is evaluate in the absence of variation (***z*** *=* (*z*,…, *z*)).

##### C.2.2 Individual fitness

Another important building block is the individual fitness *w*_*i*_ (i.e. the expected number of successful offspring) of a focal individual arbitrarily indexed *i* with trait *z*_*i*_, when its patch neighbours express the set of trait values {*z*_*j*_}_*j*≠*i*_, in a population that is otherwise monomorphic for *z*. According to our model assumptions, this individual fitness is

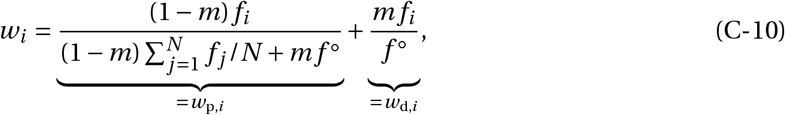

where recall *m* is the probability of dispersal;

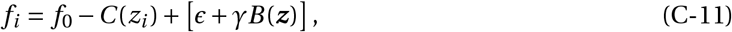

is the fecundity of the focal individual *i* (with *C* given by eq. (5), recall eq. 4 of the main text); and

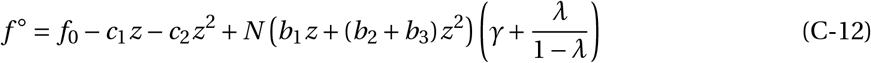

is the fecundity in a population monomorphic for *z* (obtained by substituting eqs. 2, 5 and C-7 into eq. C-11). Here and hereafter, we denote quantities evaluated under neutrality (i.e. in a population monomorphic for *z* with the environment at the corresponding ecological equilibrium 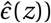 with a superscript ◦. As it will be useful later, we have decomposed individual fitness eq. (C-10) as the sum of philopatric *w*_p,*i*_ fitness (offspring that remain in their natal patch) and dispersal fitness *w*_d,*i*_ (offspring that disperse).

##### C.2.3 Relatedness

The framework found in Prigent and Mullon (2023) also depends on several relatedness coefficients that we have collected and complemented in Tables I and II.

**Appendix Table I:**
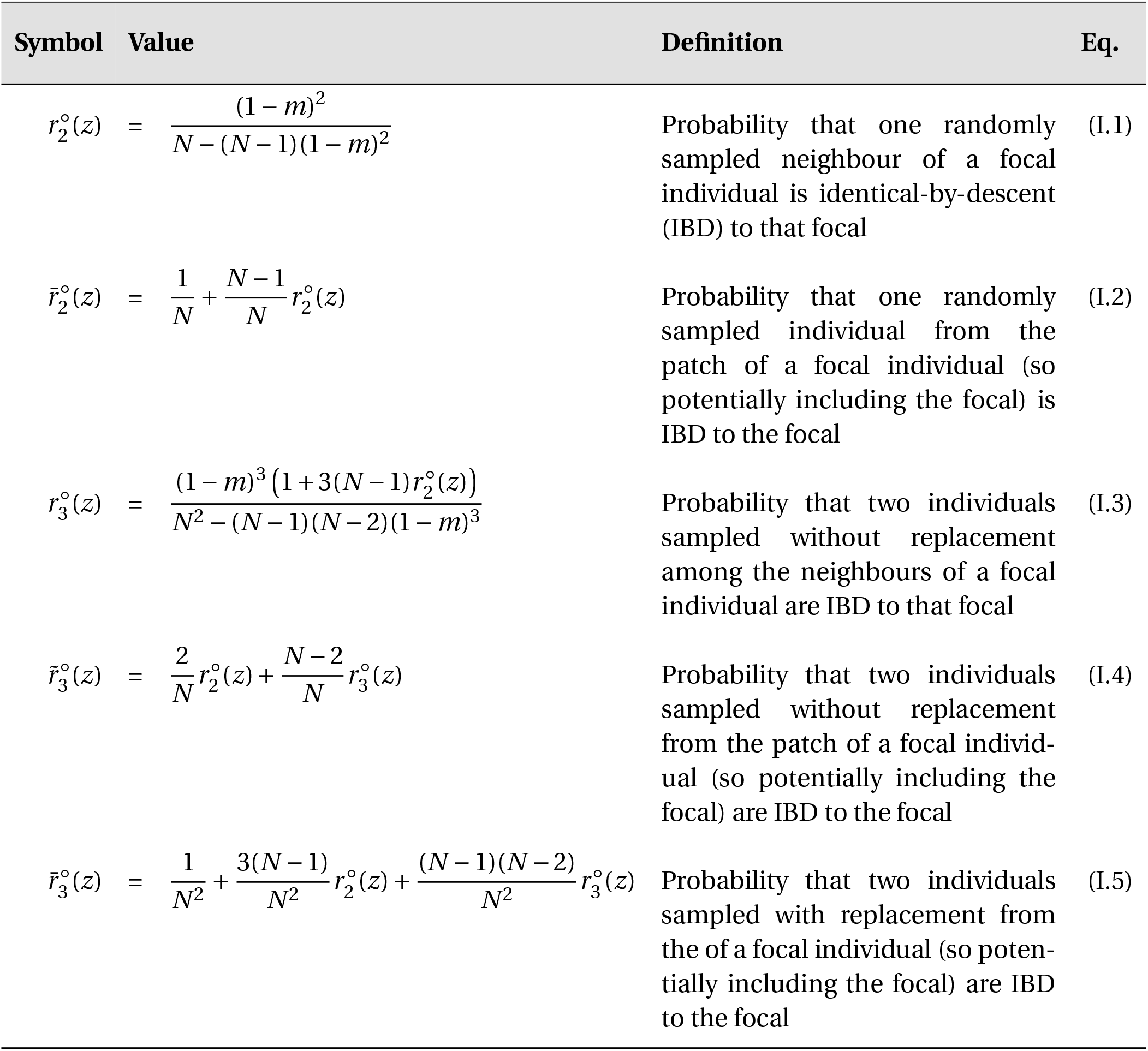
Intra-generational relatedness coefficients under neutrality necessary for invasion analyses. Recall that *N* is the (fixed) number of individuals per patch and *m* is the dispersal probability. We have written relatedness coefficients as functions of *z* to facilitate the connection with Prigent and Mullon (2023), even though here relatedness coefficients do not depend on the evolving trait.

#### C.3 Directional selection

Here we analyse the outcome of evolution under directional selection and derive eqs. 6-8 of the main text.

##### C.3.1 The selection gradient

Substituting eqs. (C-6)-(C-11) (with eqs. 2 and 5) into Prigent and Mullon (2023)’s equations (10)-(11), readily gives us the selection gradient on *z*

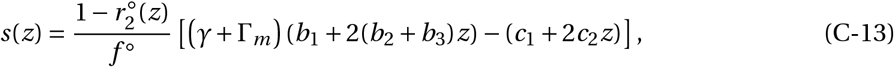

where 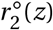 is given in eq. (I.1) in Table I and Γ_*m*_ is given by eq. (7) of the main text.

##### C.3.2 Invasion of helping

Evaluating eq. (C-13) at *z =* 0, we obtain that helping evolves when it is absent in the population if

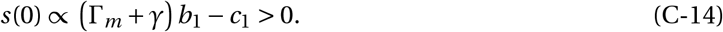

Straightforward re-arrangements of this equation yields eq. (6) of the main text, which had previously been derived by Lehmann (2007).

##### C.3.3 Convergence to evolutionary equilibrium

Using eq. (C-13) to solve *s*(*z*^*^) *=* 0 for *z*^*^ obtains the singular trait value

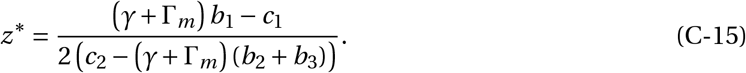

In turn, plugging eqs. (C-13) and (C-15) into eq. (C-3) we find that *z*^*^ is convergence stable when

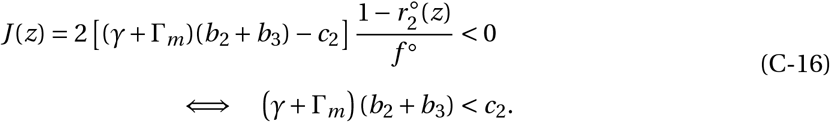

Thus, when conditions (C-14) and (C-16) are both satisfied, helping will evolve and converge to *z*^*^. When however condition (C-14) is satisfied but not (C-16), helping evolves but then selection keeps favouring ever greater values (i.e. *s*(*z*) *>* 0) until helping reaches its upper bound, *z*_max_. These considerations give us eq. (8) of the main text.

**Appendix Table II:**
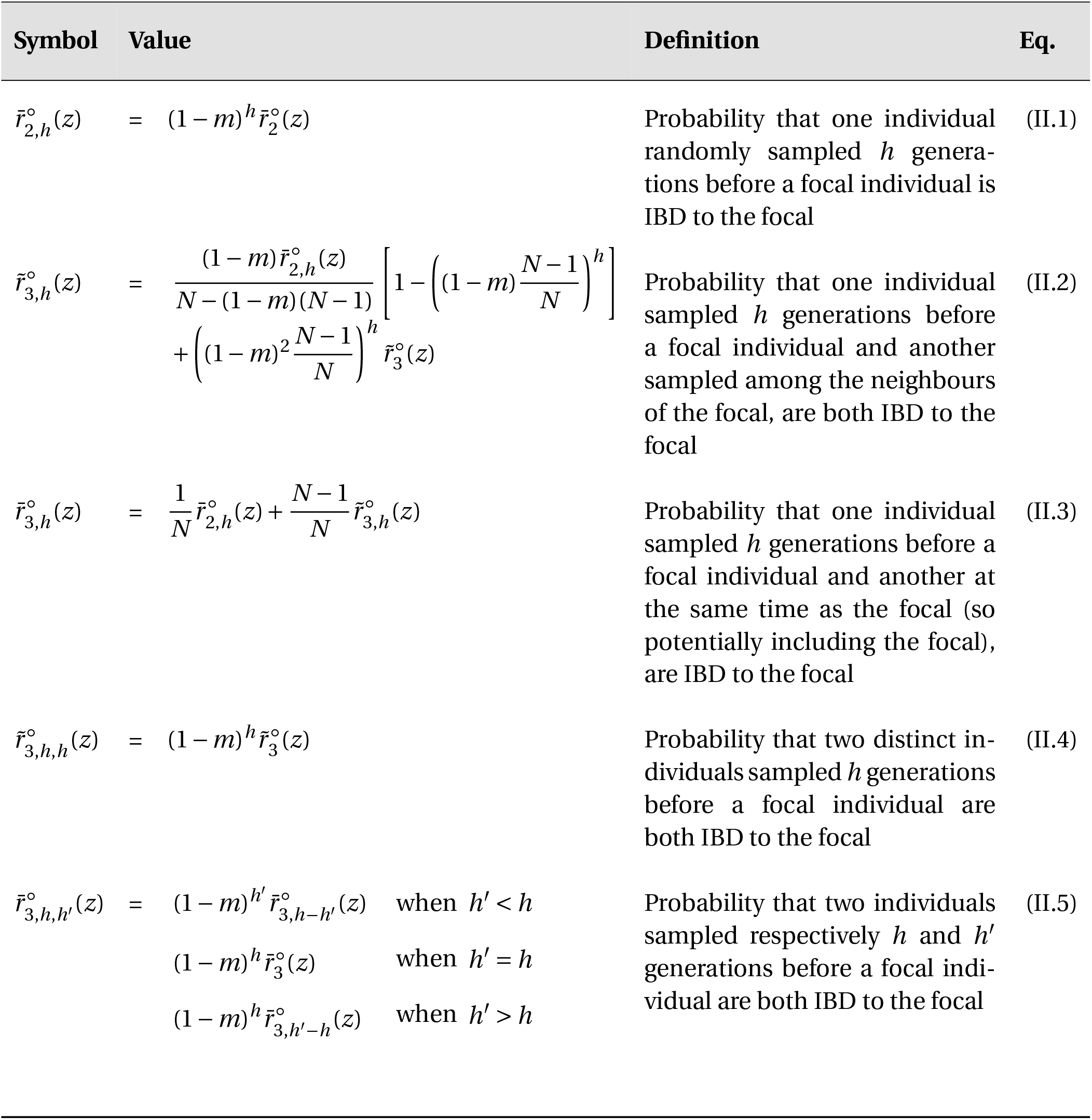
Inter-generational relatedness coefficients under neutrality necessary for invasion analyses. All these coefficients consider individuals sampled from the same patch, but at different generations. Notations are consistent with the corresponding intra-generational relatedness coefficient when 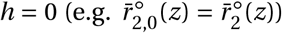. We have written relatedness coefficients as functions of *z* to facilitate the connection with Prigent and Mullon (2023), even though here relatedness coefficients do not depend on the evolving trait.

#### C.4 Disruptive selection

To obtain the coefficient of disruptive selection, we then plug eqs. (C-6)-(C-11) (with eqs. 2 and 5) into Prigent and Mullon (2023)‘s equations (15)-(22) (and using their Table 1). This gives us

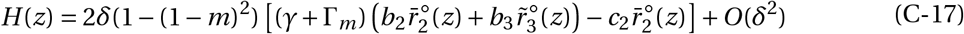

where *δ >* 0 is a parameter of the order of selection (i.e. of the order of *b*_1_, *b*_2_, *b*_3_, *c*_1_ and *c*_2_), and 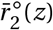 and 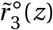 are relatedness coefficients whose expression and definitions can be found in Table I (eqs. I.2 and I.4). Straightforward re-arrangements of eq. (C-17) reveals that selection is disruptive, i.e. *H* (*z*) *>* 0, when

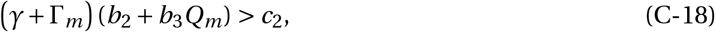

where

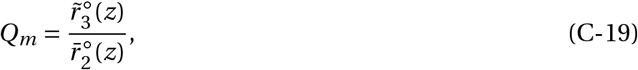

can be understood as the conditional probability that two individuals, randomly sampled without replacement from a mutant patch (i.e. a patch with at least one mutant) are both mutants, given that one of them is a mutant (as 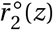 is the probability that one individual sampled from a mutant patch is a mutant). More generally, *Q*_*m*_ can be considered as a measure of the frequency of mutants in mutant patches.

#### C.5 Necessary condition for polymorphism

Next, we substitute eq. (C-16) and eq. (C-17) into eq. (C-5) to obtain,

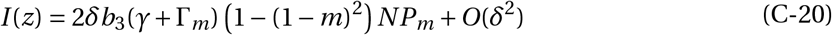

where

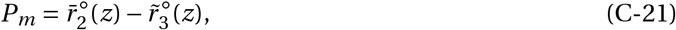

can be understood as the probability that two individuals randomly sampled without replacement from a mutant patch are not IBD (i.e. one is a mutant and another is a resident). Eq. (C-20) is eq. (10) of the main text (where we write *I = I* (*z*), as it does not depend on *z*), see there for interpretation where we dropped *δ* for ease of presentation.

#### C.6 Evolutionary branching

Putting conditions eq. (C-16) and eq. (C-18) together, we obtain that evolutionary branching occurs when

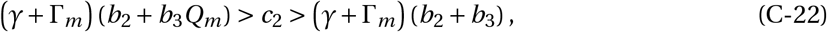

which gives eq. (12) of the main text after straightforward re-arrangements.

#### C.7 Individual-based simulations

We also ran individual-based simulations using Julia (Bezanson et al., 2017; version 1.9.2, all codes used in our manuscript can be found here : Prigent and Mullon, 2023). We simulate the evolution of a population subdivided among a large but finite number of patches, where mutations are rare with small phenotypic effect. Unless otherwise specified, the population consists of *N*_p_ *=* 2^*′*^000 patches occupied by *N =* 10 individuals. Each individual *i* ∈ {1,…, *N*} in each patch *k* ∈ {1,…, *N*_p_} is characterized by a quantitative trait *z*_*ik*_, which determines its investment into its local environment. We collect these traits values in a vector {(*z*_*ik*_)_*i,k*_} of size *N × N*_p_. Each patch is characterized by its environmental quality *ϵ*_*k*_, which we collect into a vector (*ϵ*_*k*_)_*k*_ of size *N*_p_.

We start our simulations by assuming that the population is monomorphic for an arbitrary trait value *z* (i.e. *z*_*ik*_ *= z* for all *i* and *k*), and the environment is at the ecological equilibrium for this trait value (i.e. 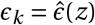 for all *k*, using eq. C-7).

We then update the collections of trait and environmental states by iterating the life-cycle described in the main text (section 2) :

During step (i), individuals invest into the environment according to eq. (2). In each patch *k*, this investment is computed as

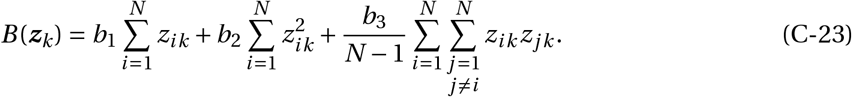

During step (ii), individuals reproduce. To model this, we calculate the fecundity *f*_*ik*_ of each individual *i* in each patch *k* according eq. (4), i.e. as

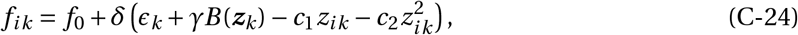

where *f*_0_ *= δ =* 1 unless stated otherwise.

Steps (iii) (migration) and (iv) (competition) are simulated in one go. We form the offspring generation by sampling *N* individuals in each patch from the parental generation, using multinomial sampling (i.e. with replacement). Each parent is weighted according to its fecundity and to the probability that its offspring will compete in the patch being filled (denoted *l*). Specifically, each parent *i* living in patch *k* is weighted by

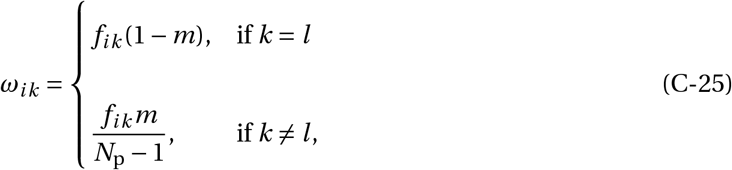

to capture random dispersal.

Each established offspring mutate with probability *µ =* 0.005, in which case we add to its trait value a value that is sampled from a Normal distribution (with parameters *µ*_z_ *=* 0, *σ*_z_ *=* 0.0025). The resulting trait value is truncated such that it remains between 0 and *z*_max_ (eq. 3).

Finally, we update the environmental variable of each patch following eq. (1), i.e. as *λ*(*ϵ*_*k*_ *+B*_*k*_ (***z***_*k*_)).

Every 1^*′*^000 generation, we sample the trait values of all individuals in the population and the environmental quality of all patches. Note that we sample the environmental quality after investment from the current generation but before environmental degradation (i.e. as *ϵ*_*k*_ *+B*_*k*_ (***z***_*k*_)), as it allows us to compare environmental quality across patches for different values of *λ* (including when *λ =* 0).

### D Fluctuating demography

Here, we detail the simulation procedure used to model fluctuating demography and obtain the results presented in section 3.3 of the main text. The procedure is broadly the same as in section C.7, except that each patch *k* is now also characterized by its size *N*_*k*_, and steps (ii-iv) (reproduction-migration-competition) are modified such that patch size *N*_*k*_ can now fluctuate between generations. Instead of sampling *N* individuals from the parental generation to model reproduction, we assume that each parent *i* from patch *k* produces a number of offspring that follows a Poisson distribution with mean *f*_*ik*_ (given by eqs. C-23 and C-24 with *N = N*_*k*_). Unless otherwise specified, we set *f*_0_ *= δ* such that *f*_*ik*_ *=* 10 when the population expresses *z = z*^*^.

Each offspring then disperses independently with probability *m* (implemented as a Bernoulli trial with probability *m* of success), in which case the patch it disperses into is sampled uniformly among the *N*_p_ − 1 remaining patches. Each offspring a patch *k* is then recruited to become an adult of the next generation with a probability 1/(1*+χN*_*k*,off_) (akin to a Beverton Holt model of regulation), where *N*_*k*,off_ is the number of offspring in the patch and *χ >* 0 is a parameter tuning the strength of density-dependent competition. To focus on the effects of evolution of fluctuations of patch size, we set *χ* such that the expected patch size *E* [*N*_*k*_] is equal to the fixed patch size *N* used previously (i.e. *E* [*N*_*k*_] *=* 10) when the population expresses *z = z*^*^.

### E Isolation-by-distance

In this appendix, we specify the analyses and simulation procedure to investigate evolution under isolation-by-distance and obtain the results presented in section 3.4 of the main text. We assume that patches are arranged according to a regular lattice over a torus to avoid boundary effects (Rousset, 2004). We consider specifically the cases of one and two-dimensional torus, and that individuals can only disperse to neighbouring patches, i.e. the stepping-stone model of dispersal. The life-cycle of the population otherwise follows the same events as under the island model of dispersal.

#### E.1 Directional selection

##### E.1.1 Selection gradient on a lattice under ecological inheritance

We first derive the selection gradient *s*(*z*) under the above assumption using previous results (Lehmann, 2008; Mullon et al., 2024) in order to study the effects of directional selection. The method is quite involved, relying on Fourier transforms and therefore we refer readers to these papers for more details, but the main idea and decomposition of directional selection follows the same line of reasoning as in the island model.

Briefly, since we assume that the local environment and the trait influences fecundity, our model falls under a specific case of the analyses found in Mullon et al. (2024). Specifically, we can use their eq. (23), where their payoff function *π* is is equal to our fecundity function (given by eq. 4). Under the assumptions made here (i.e. assuming (i) that individuals only affect and are affected by their local environment; (ii) that the environmental map only depends on the local environment; and (iii) that the number of patches is infinite), substituting their eq. (24) into their eq. (23) gives

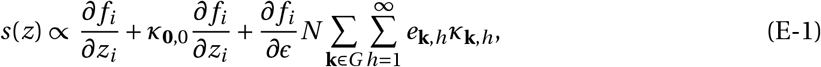

using the present notation for fecundity, where *G* is the set of possible vectorial distances among patches (i.e. the multidimensional vector of axial distances, which may be negative, p. 29 in Rousset, 2004); *κ*_**k**,*h*_ is the scaled relatedness between a focal individual and another individual living in a patch at a distance **k**, *h* generations later (eq. 19-21 of Mullon et al., 2024 for more details); and *e*_**k**,*h*_ is the effect of a change in the trait *z* of a focal individual on the environment of a patch at a distance **k**, *h* generations later (eq. 14 of Mullon et al., 2024).

Taking the limit of *κ*_**k**,*h*_ (eq. 21 in Mullon et al., 2024) when the number of patches goes to infinity gives

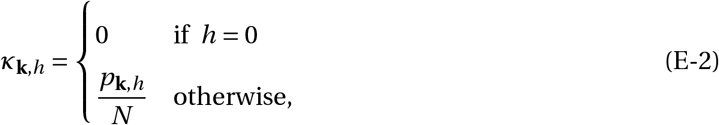

where *p*_**k**,*h*_ is the probability that under neutrality, an individual descending from a focal individual *t* generations is the future, lives in a patch at distance k from the focal. Meanwhile, since environmental modifications only affect the local patch, for *e*_**k**,*h*_ simplifies to

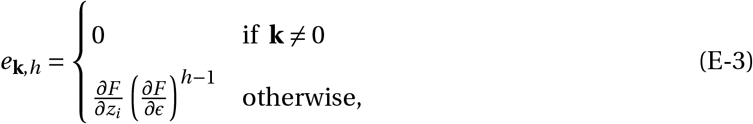

where *∂F* /*∂ϵ* is given by eq. (C-8) and *∂F* /*∂z*_*i*_ is given by eq. (C-9). Substituting eq. (E-2) and eq. (E-3) into eq. (E-1), we obtain

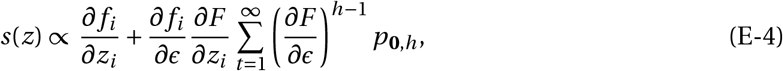

where, using eq. 22 of Mullon et al. (2024), *p*_**0**,*t*_ is given by

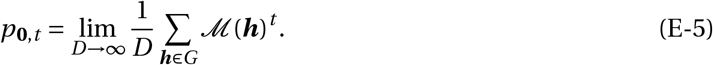

In this equation, *D* is the number of patches (and so equals the number of elements in the set G) and ℳ (***h***) is the Fourier transform of the dispersal function, i.e.

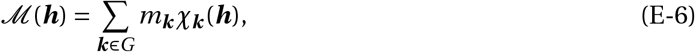

where *m*_***k***_ is the probability that an offspring disperses to a patch at distance ***k*** and

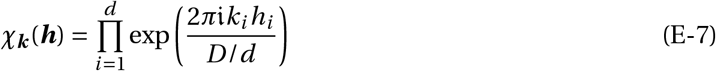

is the character function, in which 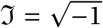, and *d* is the number of dimensions of the lattice so that *D*/*d* is the number of patches per dimension (assuming they are equally distributed across each dimension). We develop *p*_**0**,*t*_ for one- and two-dimensional lattices below.

##### E.1.2 One-dimensional lattice

Under the stepping stone model of dispersal in one dimension, the dispersal kernel is

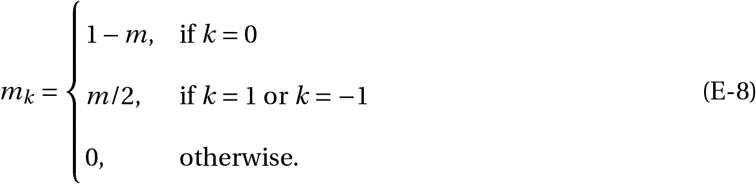

Substituting this into eq. (E-6), we obtain

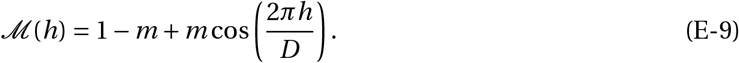

Plugging this in turn into eq. (E-5) reads as

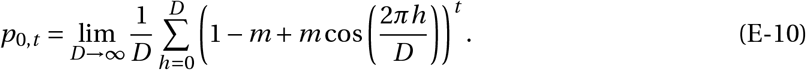

Taking the limit as the number of patches *D* → ∞, this Riemann sum converges to the integral

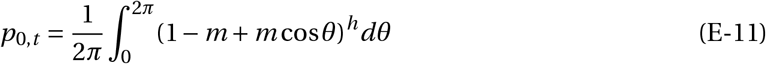

(in agreement with relatedness coefficients found in Weiss and Kimura, 1965 – their eq. 2.11 – and Duforet-Frebourg and Slatkin, 2016 – their eq. 11).

##### E.1.3 Two-dimensional lattice

In two dimensions, the dispersal kernel now is

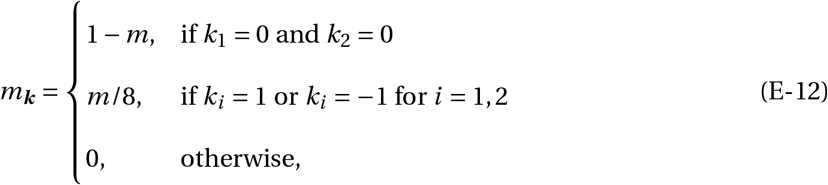

where ***k*** *=* (*k*_1_, *k*_2_), which substituted into eq. (E-6) yields

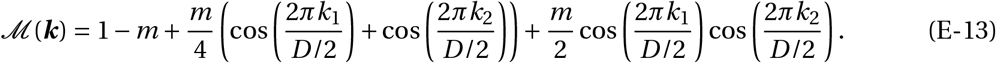

Plugging this into eq. (E-5) and taking the limit as the number of patches in each dimension goes to infinity, the resulting Riemann sum converges to the integral

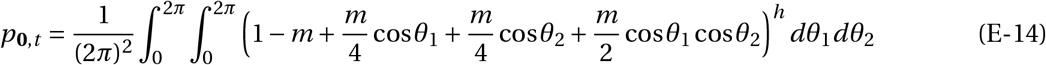

(in agreement with relatedness coefficients found in Duforet-Frebourg and Slatkin, 2016 – their eq. 12).

##### E.1.4 Putting it all together

Substituting eq. (C-8)-(C-9) and eq. (4) into eq. (E-4) we obtain

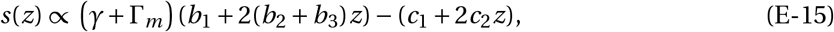

where

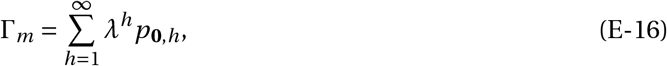

is the effect that a focal individual has on the environment of all its direct descendants.

For a one-dimensional lattice, we substitute eq. (E-11) into eq. (E-16), giving us

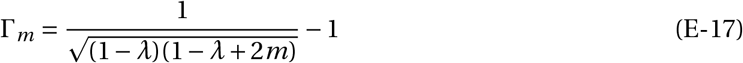

which corresponds to the first case in eq. (14) of the main text.

For a two-dimensional lattice, we substitute eq. (E-14) into eq. (E-16), giving us

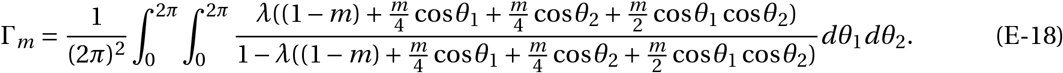

Assuming that dispersal is small, we can approximate eq. (E-18) by the second case of eq. (14) in the main text.

Solving for *s*(*z*^*^) *=* 0, we find that the evolutionary equilibrium under the stepping-stone model of dispersal has the general form

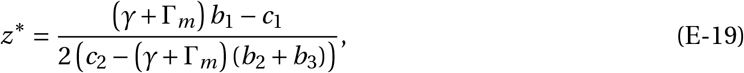

which is eq. (8) of the main text but where Γ_*m*_ is computed from the stepping stone model of dispersal (eq. E-17 for a one-dimensional model and eq. E-18 for a two-dimensional model).

#### E.2 Individual-based simulations

We used individual based simulations to investigate the emergence of polymorphism under isolation by distance. The population is distributed on a two-dimensional torus of size *N*_p_ *= N*_*x*_ *×N*_*y*_ (with *N*_x_ *= N*_y_ *=* 50, unless otherwise specified). Each patch *k* ∈ {1,…, *N*_p_} is characterized by its coordinates in the two dimensional lattice: (*x*_*k*_, *y*_*k*_), where *x*_*k*_ ∈ {1,…, *N*_x_} and *y*_*k*_ ∈ {1,…, *N*_y_}. Our algorithm is the same as the one used for the island model of dispersal (as described in Appendix C.7), except that the weights used on adults to create the next generation (eq. C-25) now incorporate the stepping stone model of dispersal – with offspring able to disperse to only one patch away from their natal patch (horizontally, vertically or diagonally, using eq. E-12). Thus, when filling patch *l*, adult *i* from patch *k* is now weighted by

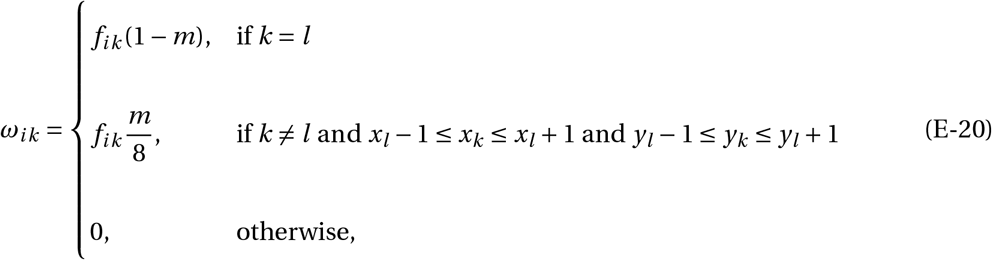

which implements the idea that an offspring has a probability *m*/8 of dispersing into each neighbouring patch and a probability 0 of dispersing into any other patch (i.e. eq. E-12).

### F Contrasts with previous results on the effects of ecological inheritance on polymorphism

Our analyses demonstrate that the combination of ecological inheritance and limited dispersal favours the emergence and maintenance of polymorphism in environmentally mediated helping. In contrast, other models have shown that ecological inheritance and limited dispersal, when considered separately, typically inhibit polymorphism. Models without ecological inheritance show that limited dispersal alone tends to suppress polymorphism by increasing kin competition (Day, 2001; Ajar, 2003; Wakano and Lehmann, 2014; Mullon and Lehmann, 2019; Parvinen et al., 2017; Schmid et al., 2024). Conversely, models without limited dispersal suggest that ecological inheritance destabilises coexistence, leading to eco-evolutionary cycles rather than a robustly stable mix of helpers and free-riders (Brown and Taddei, 2007; Weitz et al., 2016; Tilman et al., 2020).

To understand this apparent discrepancy and its biological implications, it is useful to consider that a key difference between our model and previous studies of ecological inheritance in well-mixed populations is that in these models, coexistence relies on free-riders having lower fitness than helpers in poor-quality environments (Brown and Taddei, 2007; Weitz et al., 2016; Tilman et al., 2020). This in effect assumes that an individual helper partially privatises the benefits of their investment for itself, for instance by retaining some of the produced goods before they become available to others (Smith and Schuster, 2019).

However, public goods cannot always be easily privatised. Examples include diffusible enzymes secreted by soil microbes (Allison, 2005), exopolysaccharides in bacterial biofilms (Nadell et al., 2009), and cellulolytic enzymes in termite colonies that benefit all nest members (Scharf, 2015). In plants, root exudates likewise diffuse into the rhizosphere and are accessible to both relatives and potential competitors (Badri and Vivanco, 2009). In such systems, free-riders can always access the benefits produced by helpers, making helping more vulnerable to exploitation (Rankin et al., 2007). An example of this occurs in *Pseudomonas aeruginosa*, where helping consists in the metabolically costly production of pyoverdin, a siderophore that binds and transports extracellular iron, making it available to all cells in the environment. Here, free-riders achieve their highest relative fitness when pyoverdin is rare, since they avoid production costs while still benefiting from any available pyoverdin (Kümmerli and Brown, 2010).

Instead of privatising benefits at the individual level, our model shows that ecological inheritance combined with limited dispersal allows privatisation to occur at the level of the local lineage. By structuring populations such that helpers’ descendants are more likely to inherit the benefits of past investment, this mechanism facilitates the persistence of helpers even when free-riders have a short-term advantage.

### G Other pathways for polymorphism under ecological inheritance

In this appendix, we identify other pathways via which selection can favour polymorphism under ecological inheritance by analysing trait-dependent selection (eq. C-5) for a more general model than the one described in main text section 2.1.

#### G.2 General model

We still assume the island model of dispersal as in our baseline model but now consider a general environmental map

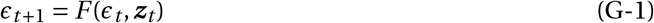

(instead of eq. C-6) with equilibrium denoted 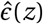, such that 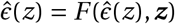 (with ***z*** *=* (*z*,…, *z*)) and 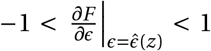, which measures the degree of ecological inheritance. Additionally, instead of eq. (C-10) (together with eq. C-11) we now consider a general individual fitness function

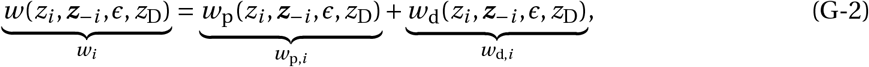

which depends on: (i) the trait of the focal individual, *z*_*i*_ ; (ii) the collection of the traits of its neighbours, ***z***_−*i*_ *=* (*z*_1_, …, *z*_*i*−1_, *z*_*i+*1_, …, *z*_*N*_) ; (iii) the state of the environment of the patch of the focal, *ϵ*; and (iv) the traits of the individuals living in all the other patches than the focal, which, for the purpose of invasion analyses, can be considered to express the same resident trait value, here denoted *z*_D_. We also decomposed fitness in terms of philopatric and dispersal components (offspring that settle in their natal and non-natal patch, respectively).

#### G.2 Trait-dependent selection

We are interested in describing *I* (*z*^*^) in terms of the general environmental map *F* and fitness function *w*_*i*_, as recall *I* (*z*^*^) *<* 0 is necessary for the emergence of polymorphism (Lehmann and Mullon, 2025). To do so, we use eq. C-5, which depends on two components: (i) the Hessian *H* (*z*^*^), which has been characterised in Prigent and Mullon (2023) (see also Ohtsuki et al., 2020); and (ii) the Jacobian 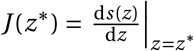 that depends on the selection gradient *s*(*z*), which has been investigated in several previous papers (e.g. Lehmann, 2007; Prigent and Mullon, 2023). We derive the Jacobian *J* (*z*^*^) from these earlier studies in Appendix G.3. Using eqs. 14-22 in Prigent and Mullon (2023) for *H* (*z*^*^) and eq. G-14a here for *J* (*z*^*^), we obtain that *I* (*z*^*^) can be decomposed as,

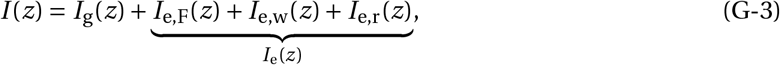

where *I*_g_(*z*) captures selection due to the intra-generational effects of trait expression, while *I*_e_(*z*) captures selection due to the inter-generational environmental effect of the trait, i.e. those due to ecological inheritance. We detail each of these below with specific interest in those due to ecological inheritance.

##### G.2.1 Trait-dependent kin selection

The first term of eq. (G-3), which is independent of ecological inheritance, is given by

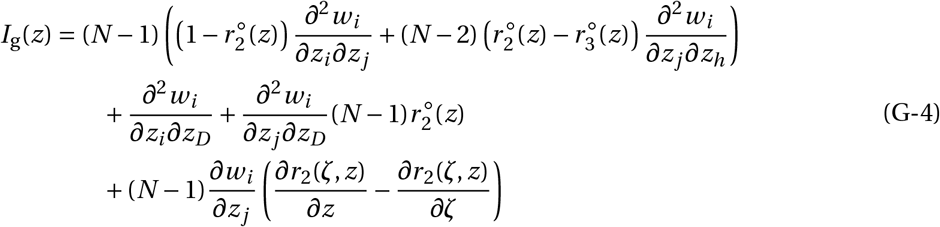

where *z* _*j*_ and *z*_*h*_ are used to denote the trait values of two different neighbours to the focal individual, and *r*_2_(*ζ, z*) is the relatedness among mutants with trait *ζ* in a resident population with trait *z* (Table III for expression of relatedness perturbations). Eq. (G-4) is equivalent to eqs. (59) and (60) of Lehmann and Mullon (2025), where it is assumed the *N =* 2 so that they are missing the term factored by (*N* − 2) on the first line of eq. (G-4). Nevertheless, the biological interpretation of eq. (G-4) is broadly the same and since we are interested here in the effects of ecological inheritance, we refer readers to Lehmann and Mullon (2025) for a detailed description of eq. (G-4).

##### G.2.2 Three main pathways for ecological inheritance to favour polymorphism

The rest of eq. (G-3), *I*_e_(*z*), is trait-dependent selection arising from ecological inheritance. Its decomposition reveals three distinct mechanisms that can favour polymorphism. For the sake of exposition and to facilitate connections with our main text, we will be considering a helping trait *z*, i.e. that is individually costly but modifies the environment in a way that increases individual fitness (which entails *∂w*_*i*_ /*∂ϵ >* 0 and *∂F* /*∂z*_*i*_ *>* 0), but our results apply more broadly. Polymorphism is favoured by ecological inheritance when *I*_e_(*z*) *<* 0. This condition implies that when residents change their trait in one direction (e.g. decrease *z*), the mutant’s best response – with respect to effects of ecological inheritance – is to change in the opposite direction (e.g. increase *z*). As a result, mutant invasion fitness increases owing to ecological inheritance as mutant and resident trait values diverge. Broadly speaking, this occurs whenever opposite trait changes in mutants and residents lead mutants, on average, to experience improved environments. There are three mechanisms via which this can happen.

###### Non-additive environmental transformations

The first mechanism by which mutants can experience improved environments when they show opposite trait changes to residents is when trait expression by different individuals has antagonistic effects on the environment. This is captured by *I*_e,F_(*z*) in eq. (G-3), which can be expressed as

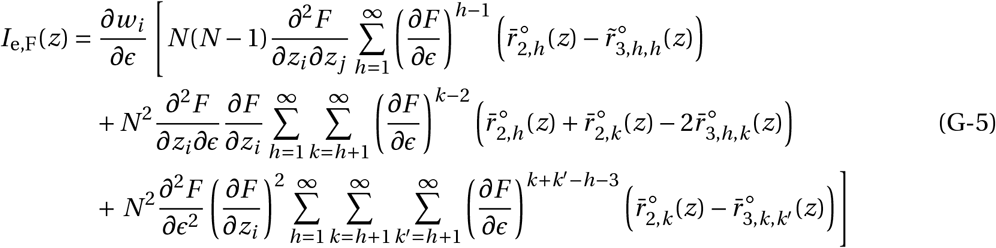

(Table II for relatedness coefficients). This equation consists of three effects that we briefly describe. Since we assume that *∂w*_*i*_ /*∂ϵ >* 0 and *∂F* /*∂z*_*i*_ *>* 0 (and all terms in parenthesis containing relatedness coefficients are positive), whether or not *I*_e,F_(*z*) is negative depends on (i) *∂*^2^*F* /(*∂z*_*i*_ *∂z*_*j*_), (ii) *∂*^2^*F* /(*∂z*_*i*_ *∂ϵ*), and (iii) *∂*^2^*F* /*∂ϵ*^2^, being negative. Effect (i) is exactly the effect we considered in the main text: antagonistic effects among individuals of the same generation on the environment (when *b*_3_ *<* 0 in eq. 2), which occurs when the effect of helping saturates with the number of helpers. Meanwhile, the remaining effects occur when, as the environment improves, (ii) the effect of trait expression on the environment shows diminishing returns, and (iii) ecological inheritance gets weaker.

Each of these effects (i)–(iii) is weighted by a relatedness coefficient that accounts for the relevant combinations of mutants and residents inhabiting the patch of a focal mutant in the present and in the past, which jointly determine the focal’s patch present environment and indirect fitness effects. For example, the coefficient 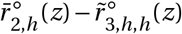, which weights the term *∂*^2^*F* /(*∂z*_*i*_ *∂z*_*j*_), is the probability that, among two individuals sampled *h* generations in the past of a focal mutant, one is mutant (i.e. related to the focal) and the other is resident (i.e. unrelated). This can be thought of as the number of times in the past that a mutant and a resident interact in the same patch and jointly modify the environment through the effect *∂*^2^*F* /(*∂z*_*i*_ *∂z*_*j*_). The others can be explained similarly.

###### Diminishing effects of the environment on individual fitness

The second mechanism that can cause mutants to see improved environments when mutants and residents show opposite trait changes is when fitness effects are reduced in improved environments. This is given by *I*_e,w_(*z*) in (eq. G-3), which can be expressed as,

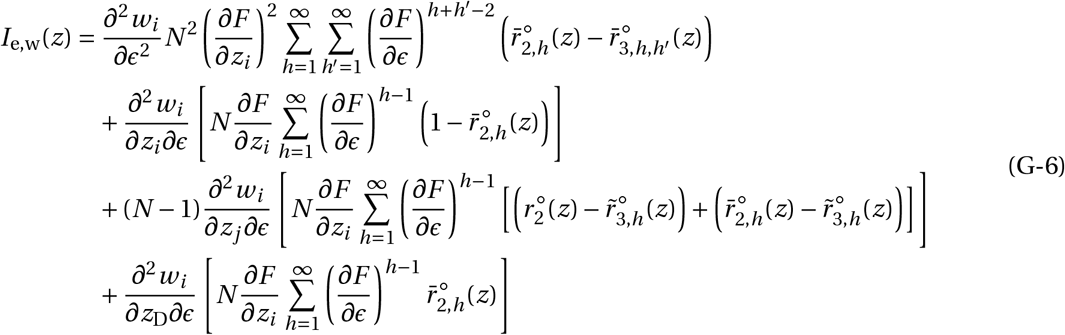

(Tables I-II for relevant relatedness coefficients). This shows that polymorphism is favoured, when as the environment improves (i.e. as *ϵ* increases), the fitness effects of the environment (*∂*^2^*w*_*i*_ /*∂ϵ*^2^ *<* 0), of the focal’s trait (*∂*^2^*w*_*i*_ /(*∂z*_*i*_ *∂ϵ*) *<* 0), of patch neighbours’ traits (*∂*^2^*w*_*i*_ /(*∂z*_*j*_ *∂ϵ*) *<* 0), and of the trait of the individuals in other patches (*∂*^2^*w*_*i*_ /(*∂z*_D_*∂ϵ*) *<* 0), diminish. Again, each of these effects in eq. (G-6) is weighted by a relatedness coefficient that accounts for the relevant combinations of mutants and residents inhabiting the patch of a focal mutant in the present and in the past.

###### Inter-generational relatedness effects

Finally, the third and final mechanism for ecological inheritance to promote polymorphism is when opposite trait changes in mutants and residents make mutants more likely to inherit environments improved by related ancestors. This is captured by the last component of eq. (G-3),

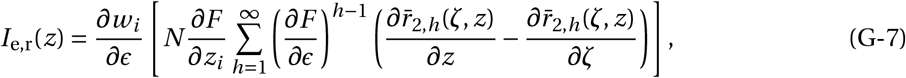

which shows that *I*_e,r_(*z*) *<* 0 requires that 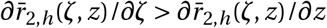 (under our assumptions that *∂w*_*i*_ /*∂ϵ >* 0 and *∂F* /*∂z*_*i*_ *>* 0). This condition says that the effect of a trait change in a mutant lineage on its own inter-generational relatedness is greater than the effect of a corresponding change in the resident population. When this holds, opposite trait changes in mutants and residents entail that a mutant is more likely to experience the improved environmental conditions that its ancestors have created. This could for instance occur where environmentally mediated helping is negatively associated with dispersal or patch extinction.

These three pathways—non-additive environmental interactions, diminishing fitness returns from environmental improvements, and inter-generational relatedness effects—collectively highlight how ecological inheritance and spatial structure can jointly promote and stabilise adaptive polymorphisms.

#### G.3 Computing the Jacobian

In this section, we compute the Jacobian *J* (*z*) from the selection gradient *s*(*z*) using eq. (C-3). The selection gradient for our model is given by eq. (10) in Prigent and Mullon (2023), i.e.

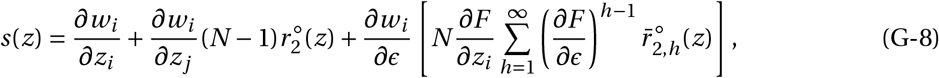

where relatedness coefficients are described in Tables I and II. Let us now substitute eq. (G-8) into eq. (C-3). Using the product rule, we obtain

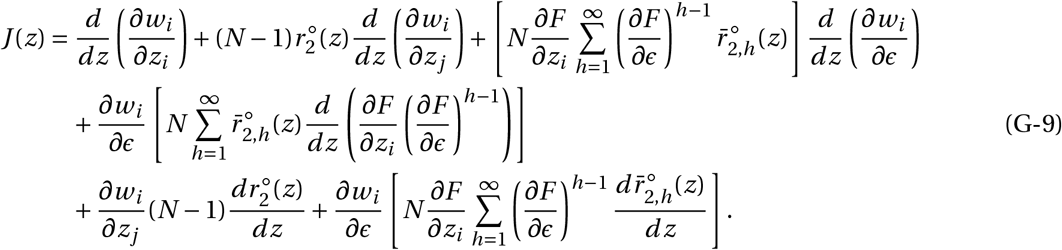

This depends on the total derivatives of several quantities with respect to *z* (i.e. *d* /*dz*), which we develop below making repeated use of the chain rule.

First, from the fitness function eq. (G-2), we readily obtain

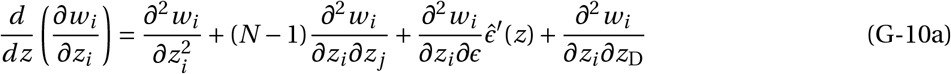

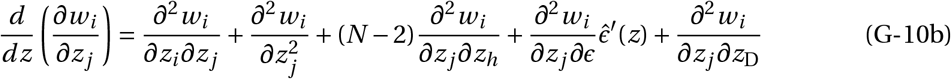

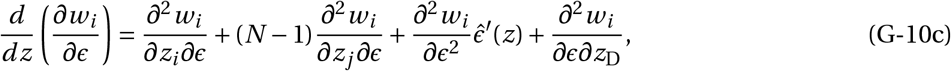

where recall that each partial derivative is estimated in a monomorphic population for *z* and where 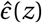 is at the associated equilibrium, i.e. such that 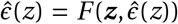 with *z*_*i*_ *= z* for all *i*. Taking the derivative of both sides of the equation 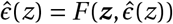 with respect to *z* and solving for 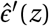 yields

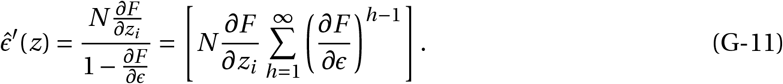

We also use the product rule to obtain

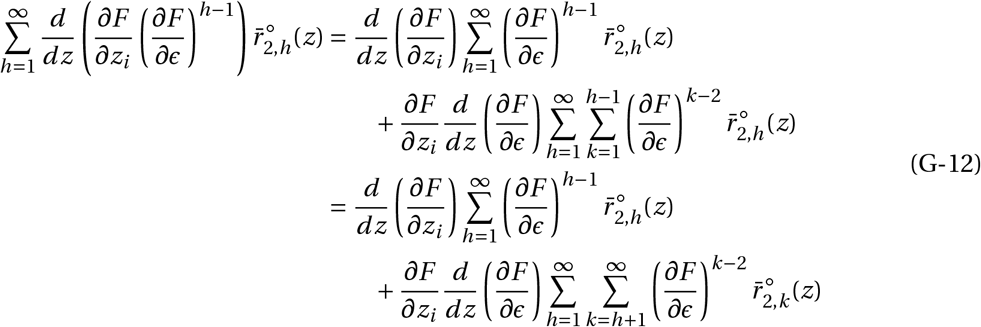

where we operated a change in indices in the sums to go from the first to the second equality. Meanwhile, using eqs. (G-1) and (G-11), we find

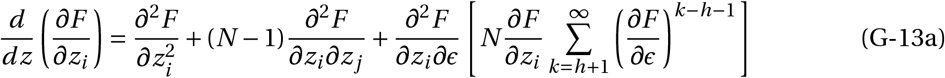

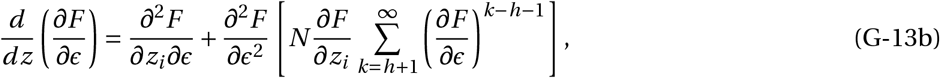

where the term between square brackets is equal to 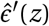 (eq. G-11 with a change of indices to facilitate later interpretation).

We obtain *J* (*z*) by substituting eqs. (G-13) into eq. (G-12), then eq. (G-12) and eqs. (G-10) with eq. (G-11) into eq. (G-9), which after re-arrangements gives

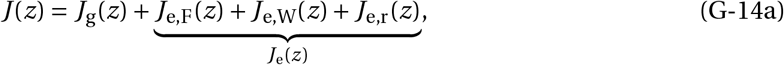

where

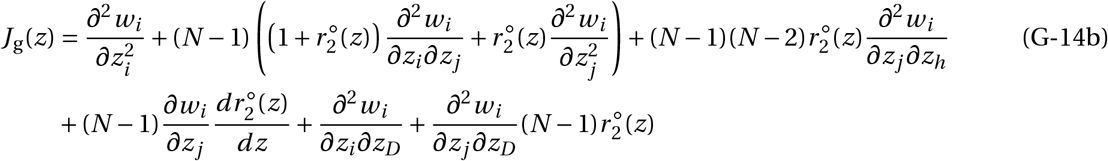

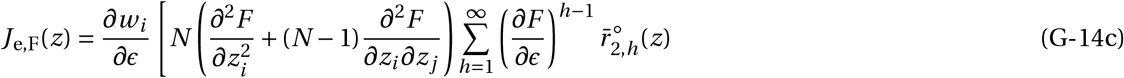

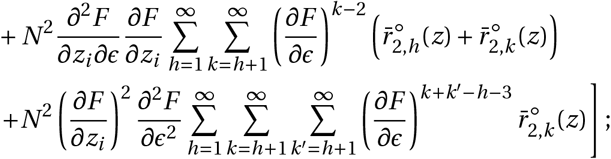

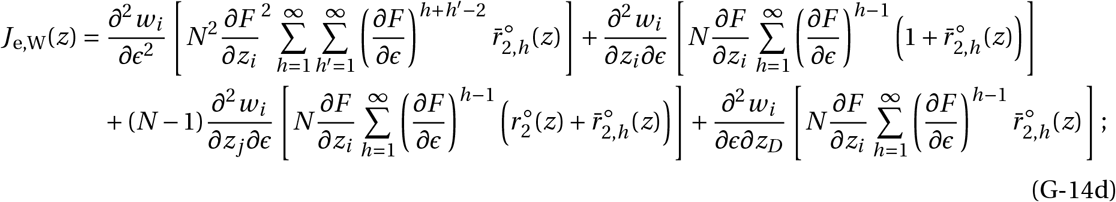

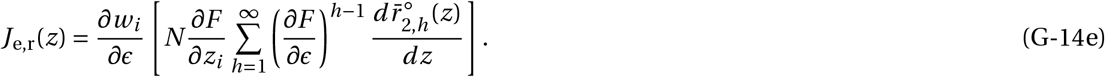

The expressions of 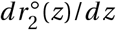 and 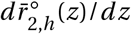 in terms of fitness effects can be found in Table III (eq. III.3-III.6). We detail their derivation in section G.4)

#### G.4 Effects of trait on relatedness coefficients

We derive here 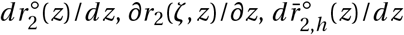and 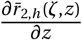, which appear in *J* (*z*) and therefore *I* (*z*). We do not make use of these in the present paper but derive them for the sake of completeness, should readers be interested in making of the framework.

##### G.4.1 Intra-generational relatedness

Let us first re-write 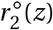 (eq. I.1) as

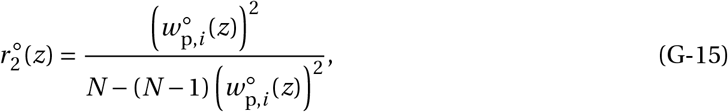

where 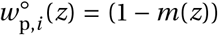 is the philopatric component of fitness under neutrality (eq. G-2, where *m*(*z*) is the backward probability of dispersal). Taking the derivative of eq. (G-15), we obtain

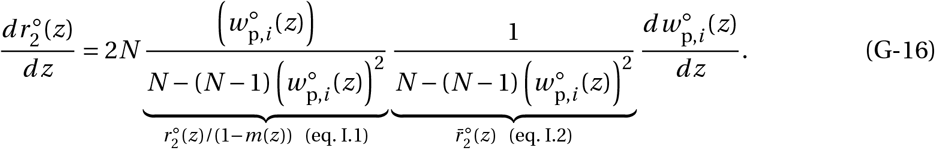

We can derive this last derivative from eq. (G-2) using the chain rule, obtaining

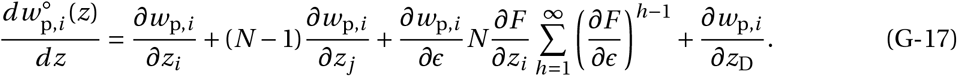

Substituting this into eq. (G-16), we obtain

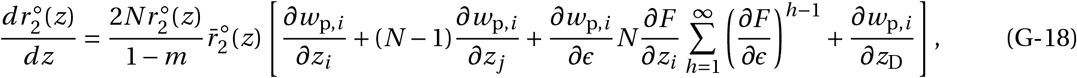

as required.

We now compute *∂r*_2_(*ζ, z*)/*∂z*. To do so, note first that by definition, we have

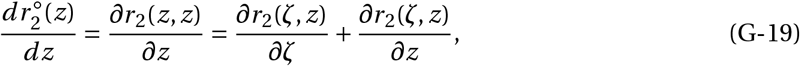

which can be re-arranged into

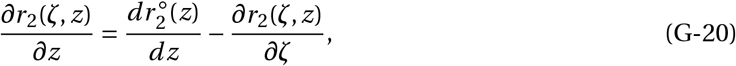

in which we substitute the value of 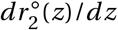 obtained in eq. (G-18) and the value of *∂r*_2_(*ζ, z*)/*∂ζ* from eq. (III.1) (eq. E-22 in Prigent and Mullon, 2023, see their Appendix E for derivation), obtaining

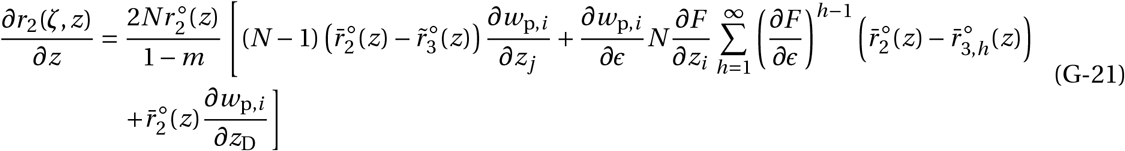

(Tables I-II for values of the relevant relatedness coefficients).

##### G.4.2 Inter-generational relatedness

We proceed similarly to the above section. We first rewrite 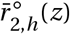 (eq. II.1) as

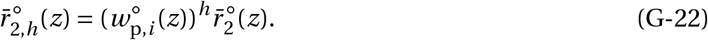

Taking the derivative of eq. (G-22) with respect to *z*, we obtain

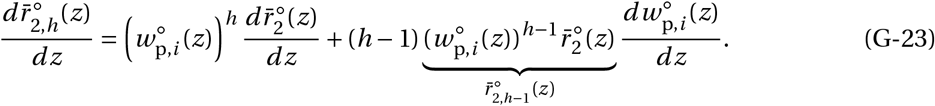

Using eq. (I.2), the derivative of 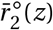 is simply

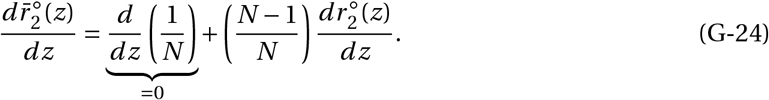

Substituting eq. (G-24) and eq. (G-17) into eq. (G-23), we obtain

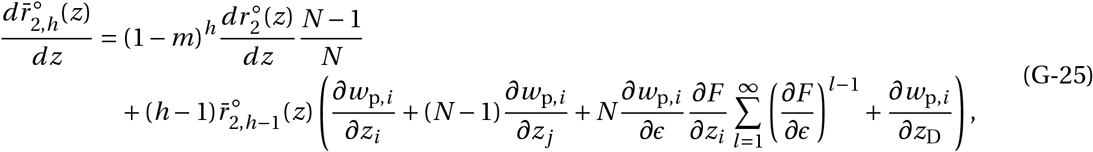

which can be expressed as

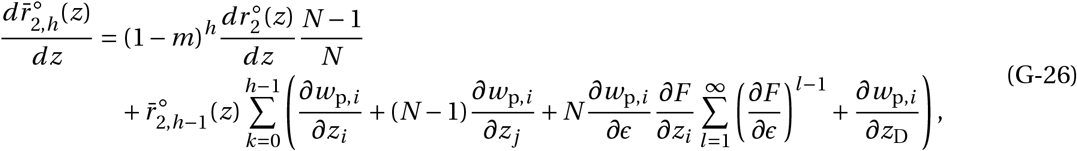

which turns out to be more useful.

Finally, we turn to 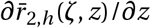, which can be decomposed into

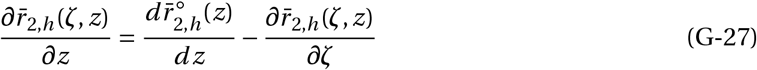

in which we substitute 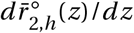 from eq. (G-26) and 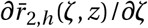 from eq. (III.4) (from eq. E-48 in Prigent and Mullon, 2023, see their Appendix E for derivation), obtaining

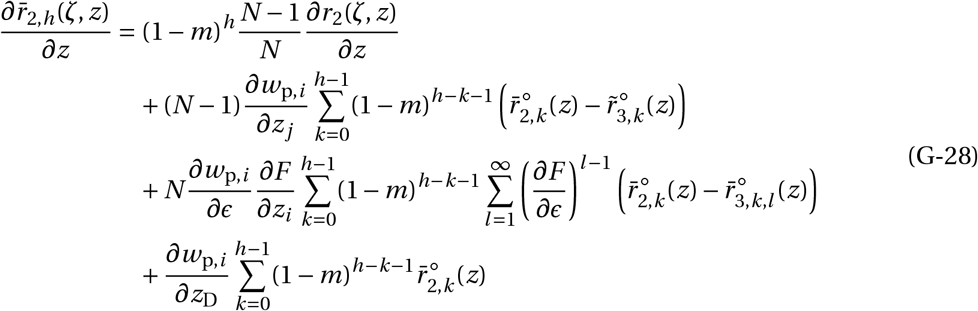

(Table II for values of the relevant relatedness coefficients).

**Appendix Table III:**
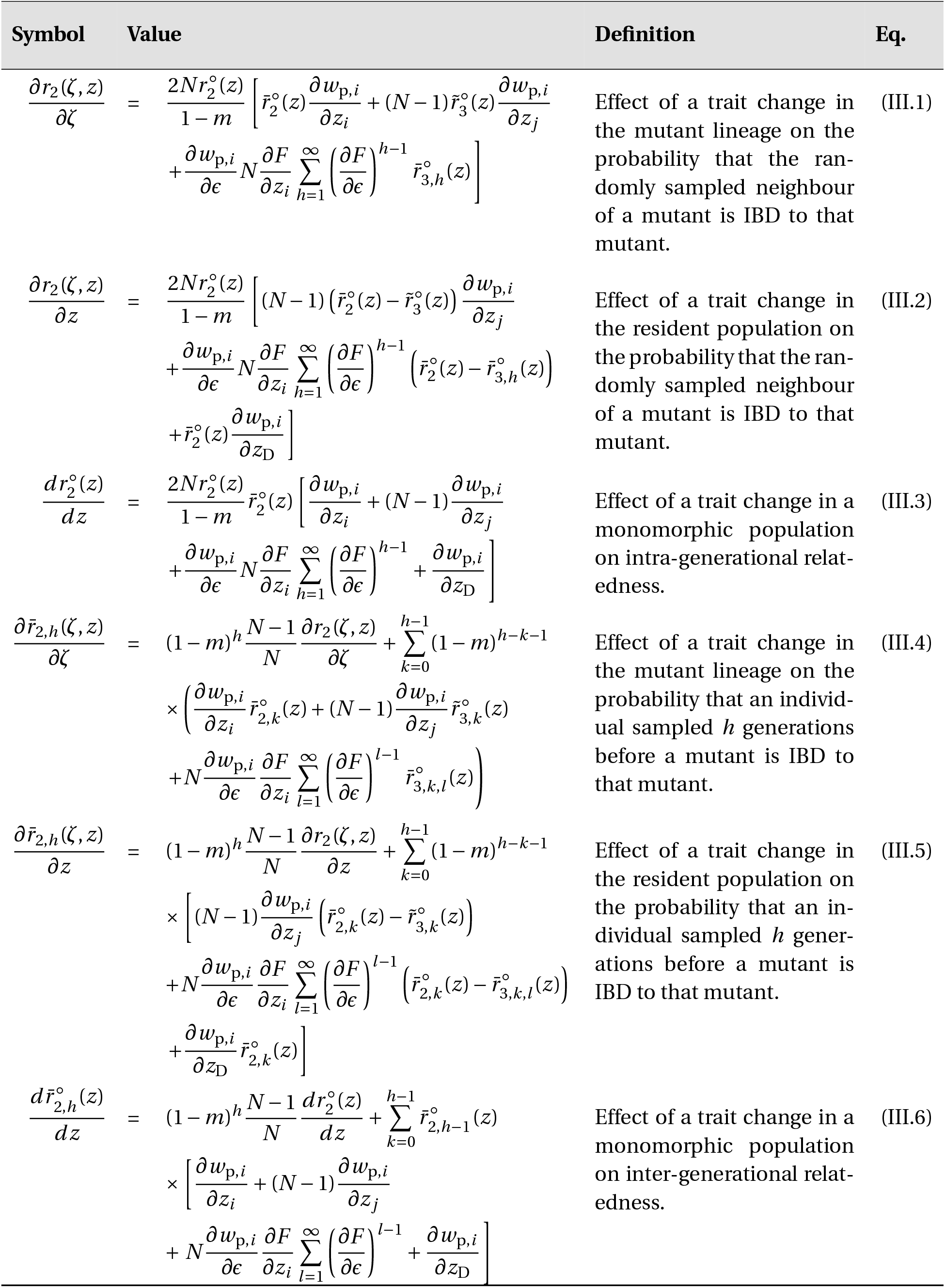
Effects of a trait change on intra- and inter-generational relatedness relevant for the analyses of selection. Eq (III.1) and (III.4) are derived in Prigent and Mullon (2023) Appendix E, the other equations in Appendix G.4. These are shown for the sake of completeness.

## Notes

### Competing Interest Statement

The authors have declared no competing interest.

### Funding

- Swiss National Science Foundation: PCEFP3_181243

## References

Ajar, É. 2003. Analysis of disruptive selection in subdivided populations. BMC Evolutionary Biology 3:1–12.

Allen, B., J. Gore, and M. A. Nowak. 2013. Spatial dilemmas of diffusible public goods. Elife 2:e01169.

Allison, S. D. 2005. Cheaters, diffusion and nutrients limit the optimization of enzyme production by microbes. Ecology Letters 8:626–635.

Avila, P. and C. Mullon. 2023. Evolutionary game theory and the adaptive dynamics approach: adaptation where individuals interact. Philosophical Transactions of the Royal Society B 378:20210502.

Ayala, F. J. and C. A. Campbell. 1974. Frequency-dependent selection. Annual review of Ecology and systematics pp. 115–138.

Badri, D. V. and J. M. Vivanco. 2009. Regulation and function of root exudates. Plant, Cell & Environment 32:666–681.

Bayard, B. and C. Jolly. 2007. Environmental behavior structure and socio-economic conditions of hillside farmers: A multiple-group structural equation modeling approach. Ecological economics 62:433–440.

Bever, J. D. 1994. Feedback between plants and their soil communities in an old field community. Ecology 75:1965–1977.

Bever, J. D. 2002. Negative feedback within a mutualism: host-specific growth of mycorrhizal fungi reduces plant benefit. Proceedings of the Royal Society B: Biological Sciences 269:2595–2601.

Bezanson, J., A. Edelman, S. Karpinski, and V. B. Shah. 2017. Julia: A fresh approach to numerical computing. SIAM review 59:65–98.

Bolnick, D. I., P. Amarasekare, M. S. Araújo, R. Bürger, J. M. Levine, M. Novak, V. H. Rudolf, S. J. Schreiber, M. C. Urban, and D. A. Vasseur. 2011. Why intraspecific trait variation matters in community ecology. Trends in ecology & evolution 26:183–192.

Bolnick, D. I., R. Svanbäck, J. A. Fordyce, L. H. Yang, J. M. Davis, C. D. Hulsey, and M. L. Forister. 2003. The ecology of individuals: incidence and implications of individual specialization. The American Naturalist 161:1–28.

Borenstein, D. B., Y. Meir, J. W. Shaevitz, and N. S. Wingreen. 2013. Non-local interaction via diffusible resource prevents coexistence of cooperators and cheaters in a lattice model. PloS one 8:e63304.

Bouchard, T. J. 2004. Genetic influence on human psychological traits: A survey. Current Directions in Psychological Science 13:148–151.

Brown, S. P. and F. Taddei. 2007. The durability of public goods changes the dynamics and nature of social dilemmas. PLoS One 2:e593.

Calderone, N. W. and R. E. Page. 1988. Genotypic variability in age polyethism and task specialization in the honey bee, apis mellifera (hymenoptera: Apidae). Behavioral ecology and sociobiology 22:17– 25.

Carmi, N. 2013. Caring about tomorrow: Future orientation, environmental attitudes and behaviors. Environmental Education Research 19:430–444.

Cavalli-Sforza, L. L. and M. W. Feldman. 1978. Darwinian selection and “altruism”. Theoretical population biology 14:268–280.

Clarke, B. and P. O’Donald. 1964. Frequency-dependent selection. Heredity pp. 201—-206.

Clobert, J. 2012. Dispersal ecology and evolution. Oxford University Press.

Clutton-Brock, T., A. Russell, and L. Sharpe. 2003. Meerkat helpers do not specialize in particular activities. Animal Behaviour 66:531–540.

Clutton-Brock, T. H. and G. A. Parker. 1995. Punishment in animal societies. Nature 373:209–216.

Crespi, B. J. 2001. The evolution of social behavior in microorganisms. Trends in ecology & evolution 16:178–183.

Dawkins, R. 1982. The Extended Phenotype. Oxford University Press, Oxford.

Dawkins, R. 2004. Extended phenotype–but not too extended. a reply to laland, turner and jablonka. Biology and Philosophy 19:377–396.

Day, T. 2001. Population structure inhibits evolutionary diversification under competition for resources. Microevolution Rate, Pattern, Process pp. 71–86.

Dercole, F. and S. Rinaldi. 2008. Analysis of evolutionary processes: the adaptive dynamics approach and its applications. Princeton University Press.

Doebeli, M. and U. Dieckmann. 2000. Evolutionary branching and sympatric speciation caused by different types of ecological interactions. The american naturalist 156:S77–S101.

Doebeli, M. and C. Hauert. 2005. Models of cooperation based on the prisoner’s dilemma and the snowdrift game. Ecology letters 8:748–766.

Doebeli, M., C. Hauert, and T. Killingback. 2004. The evolutionary origin of cooperators and defectors. science 306:859–862.

Dreller, C., M. Fondrk, and R. Page. 1995. Genetic variability affects the behavior of foragers in a feral honeybee colony. Naturwissenschaften 82:243–245.

Duforet-Frebourg, N. and M. Slatkin. 2016. Isolation-by-distance-and-time in a stepping-stone model. Theoretical population biology 108:24–35.

Foster, K. R. 2004. Diminishing returns in social evolution: the not-so-tragic commons. Journal of Evolutionary Biology 17:1058–1072.

Geritz, S. A., E. Kisdi, G. Meszé NA, and J. A. Metz. 1998. Evolutionarily singular strategies and the adaptive growth and branching of the evolutionary tree. Evolutionary ecology 12:35–57.

Gerlee, P. and P. M. Altrock. 2019. Persistence of cooperation in diffusive public goods games. Physical Review E 99:062412.

Grinsted, L. and J. Field. 2018. Predictors of nest growth: diminishing returns for subordinates in the paper wasp polistes dominula. Behavioral Ecology and Sociobiology 72:1–8.

Heino, M., A. J. Metz, and V. Kaitala. 1998. The enigma of frequency-dependent selection. TREE 13:367–370.

Hirsh, J. B. 2010. Personality and environmental concern. Journal of Environmental Psychology 30:245–248.

Houslay, T. M., J. F. Nielsen, and T. H. Clutton-Brock. 2021. Contributions of genetic and nongenetic sources to variation in cooperative behavior in a cooperative mammal. Evolution 75:3071–3086.

Ito, H. and M. Yamamichi. 2024. A complete classification of evolutionary games with environmental feedback. PNAS nexus 3:pgae455.

Jeanson, R. and A. Weidenmüller. 2014. Interindividual variability in social insects–proximate causes and ultimate consequences. Biological Reviews 89:671–687.

Julian, G. E. and J. H. Fewell. 2004. Genetic variation and task specialization in the desert leaf-cutter ant, acromyrmex versicolor. Animal Behaviour 68:1–8.

Kandler, C. 2012. Nature and nurture in personality development: The case of neuroticism and extraversion. Current Directions in Psychological Science 21:290–296.

Kimura, M. 1953. “stepping-stone” model of population. Ann. Rep. Nat. Inst. Genet. Japan pp. 62–63.

Kisdi, E. and S. A. H. Geritz. 2010. Adaptive dynamics: A framework to model evolution in the ecological theatre. Journal of Mathematical Biology 61:165–169.

Koenig, W. D. and J. L. Dickinson. 2004. Ecology and evolution of cooperative breeding in birds. Cambridge University Press.

Komdeur, J. 2006. Variation in individual investment strategies among social animals. Ethology 112:729–747.

Kümmerli, R. and S. P. Brown. 2010. Molecular and regulatory properties of a public good shape the evolution of cooperation. Proceedings of the National Academy of Sciences 107:18921–18926.

Laland, K. N., F. J. Odling-Smee, and M. W. Feldman. 1996. The evolutionary consequences of niche construction: a theoretical investigation using two-locus theory. Journal of evolutionary biology 9:293–316.

Laland, K. N., F. J. Odling-Smee, and M. W. Feldman. 1999. Evolutionary consequences of niche construction and their implications for ecology. Proceedings of the National Academy of Sciences 96:10242–10247.

Lehmann, L. 2007. The evolution of trans-generational altruism: kin selection meets niche construction. Journal of Evolutionary Biology 20:181–189.

Lehmann, L. 2008. The adaptive dynamics of niche constructing traits in spatially subdivided populations: evolving posthumous extended phenotypes. Evolution: International Journal of Organic Evolution 62:549–566.

Lehmann, L. and C. Mullon. 2025. Evolution of quantitative traits: exploring the ecological, social and genetic bases of adaptive polymorphism. bioRxiv.

Leimar, O. and J. M. McNamara. 2023. Game theory in biology: 50 years and onwards. Philosophical Transactions of the Royal Society B 378:20210509.

Lewontin, R. C. 1958. A general method for investigating the equilibrium of gene frequency in a population. Genetics 43:419.

MacLean, R. C., A. Fuentes-Hernandez, D. Greig, L. D. Hurst, and I. Gudelj. 2010. A mixture of “cheats” and “co-operators” can enable maximal group benefit. PLoS biology 8:e1000486.

Maynard Smith, J. 1962. Disruptive selection, polymorphism and sympatric speciation. Nature.

Maynard Smith, J. 1982. Evolution and the Theory of Games. Cambridge University Press.

Maynard Smith, J. and G. R. Price. 1973. The logic of animal conflict. Nature 246:15–18.

Milfont, T. L. and V. V. Gouveia. 2006. Time perspective and values: An exploratory study of their relations to environmental attitudes. Journal of environmental psychology 26:72–82.

Mullon, C. and L. Lehmann. 2018. Eco-evolutionary dynamics in metacommunities: ecological inheritance, helping within species, and harming between species. The American Naturalist 192:664– 686.

Mullon, C. and L. Lehmann. 2019. An evolutionary quantitative genetics model for phenotypic (co) variances under limited dispersal, with an application to socially synergistic traits. Evolution 73:1695–1728.

Mullon, C., J. Peña, and L. Lehmann. 2024. The evolution of environmentally mediated social interactions and posthumous spite under isolation by distance. PLoS computational biology 20:e1012071.

Mullon, C., J. Y. Wakano, and H. Ohtsuki. 2021. Coevolutionary dynamics of genetic traits and their long-term extended effects under non-random interactions. Journal of Theoretical Biology 525:110750.

Nadell, C. D., J. B. Xavier, and K. R. Foster. 2009. The sociobiology of biofilms. FEMS Microbiology Reviews 33:206–224.

Nichols, H. J., K. Arbuckle, J. L. Sanderson, E. I. Vitikainen, H. H. Marshall, F. J. Thompson, M. A. Cant, and D. A. Wells. 2021. A double pedigree reveals genetic but not cultural inheritance of cooperative personalities in wild banded mongooses. Ecology Letters 24:1966–1975.

Odling-Smee, F. J. 1988. Niche-constructing phenotypes. In The role of behavior in evolution. The MIT Press.

Odling-Smee, F. J., K. N. Laland, and M. W. Feldman. 2003. Niche Construction: The Neglected Process in Evolution. Princeton University Press.

Ohtsuki, H., C. Rueffler, J. Y. Wakano, K. Parvinen, and L. Lehmann. 2020. The components of directional and disruptive selection in heterogeneous group-structured populations. Journal of Theoretical Biology 507:110449.

O’Donnell, S. 1996. Rapd markers suggest genotypic effects on forager specialization in a eusocial wasp. Behavioral ecology and sociobiology 38:83–88.

Page, R. E. and G. E. Robinson. 1991. The genetics of division of labour in honey bee colonies. In Advances in insect physiology, vol. 23, pp. 117–169. Elsevier.

Parvinen, K., H. Ohtsuki, and J. Y. Wakano. 2017. The effect of fecundity derivatives on the condition of evolutionary branching in spatial models. Journal of Theoretical Biology 416:129–143.

Pontarotti, G. 2022. Environmental inheritance: Conceptual ambiguities and theoretical issues. Biological Theory 17:36–51.

Prigent, I. and C. Mullon. 2023. The molding of intraspecific trait variation by selection under ecological inheritance. Evolution 77:2144–2161.

Prigent, I. and C. Mullon. 2025. Online supplementary material for “ecological inheritance facilitates the coexistence of environmental helpers and free-riders”. https://github.com/iris-prigent/Supplementary_Ecological_inheritance_facilitates_coexistence.

Rankin, D. J., K. Bargum, and H. Kokko. 2007. The tragedy of the commons in evolutionary biology. Trends in ecology & evolution 22:643–651.

Rapoport, A. and A. M. Chammah. 1965. Prisoner’s dilemma: A study in conflict and cooperation, vol. 165. University of Michigan press.

Rickinson, M. 2001. Learners and learning in environmental education: A critical review of the evidence. Environmental education research 7:207–320.

Riehl, C. and M. E. Frederickson. 2016. Cheating and punishment in cooperative animal societies. Philosophical Transactions of the Royal Society B: Biological Sciences 371:20150090.

Robinson, G. 1989. Genetic basis for division of labor in an insect society. The Genetics of Social Evolution pp. 61–80.

Rousset, F. 2004. Genetic Structure and Selection in Subdivided Populations, vol. 40. Princeton University Press.

Rousset, F. and O. Ronce. 2004. Inclusive fitness for traits affecting metapopulation demography. Theoretical population biology 65:127–141.

Rueffler, C., T. J. Van Dooren, and J. A. Metz. 2006. The evolution of resource specialization through frequency-dependent and frequency-independent mechanisms. The American Naturalist 167:81– 93.

Ruel, J. J. and M. P. Ayres. 1999. Jensen’s inequality predicts effects of environmental variation. Trends in Ecology & Evolution 14:361–366.

Ruzicka, F., M. K. Zwoinska, D. Goedert, H. Kokko, X.-Y. L. Richter, I. R. Moodie, S. Nilén, C. Olito, E. I. Svensson, P. Czuppon, and T. Connallon. 2025. A century of theories of balancing selection. bioRxiv.

Scharf, M. E. 2015. Termites as targets and models for biotechnology. Annual Review of Entomology 60:77–102.

Scheuring, I. 2014. Diffusive public goods and coexistence of cooperators and cheaters on a 1d lattice. Plos one 9:e100769.

Schmid, M., C. Rueffler, L. Lehmann, and C. Mullon. 2024. Resource variation within and between patches: Where exploitation competition, local adaptation, and kin selection meet. The American Naturalist 203:E19–E34.

Sibly, R. M. and R. Curnow. 2011. Selfishness and altruism can coexist when help is subject to diminishing returns. Heredity 107:167–173.

Siegmann, S., R. Feitsch, D. W. Hart, N. C. Bennett, D. J. Penn, and M. Zöttl. 2021. Naked mole-rats (heterocephalus glaber) do not specialise in cooperative tasks. Ethology 127:850–864.

Silver, M. and E. Di Paolo. 2006. Spatial effects favour the evolution of niche construction. Theoretical Population Biology 70:387–400.

Skulason, S. and T. B. Smith. 1995. Resource polymorphisms in vertebrates. Trends in ecology & evolution 10:366–370.

Slatkin, M. 1980. Ecological character displacement. Ecology 61:163–177.

Smith, P. and M. Schuster. 2019. Public goods and cheating in microbes. Current biology 29:R442– R447.

Smith, T. B. and S. Skúlason. 1996. Evolutionary significance of resource polymorphisms in fishes, amphibians, and birds. Annual review of ecology and systematics 27:111–133.

Sozou, P. D. 2009. Individual and social discounting in a viscous population. Proceedings of the Royal Society B: Biological Sciences 276:2955–2962.

Taylor, P. D. 1992. Altruism in viscous populations—an inclusive fitness model. Evolutionary ecology 6:352–356.

Thorley, J., R. Mendonça, P. Vullioud, M. Torrents-Ticó, M. Zöttl, D. Gaynor, and T. Clutton-Brock. 2018. No task specialization among helpers in damaraland mole-rats. Animal Behaviour 143:9–24.

Tilman, A. R., J. B. Plotkin, and E. Akçay. 2020. Evolutionary games with environmental feedbacks. Nature communications 11:915.

Trivers, R. 1985. Social evolution. Benjamin-Cummings Pub Co.

Van der Putten, W. H., M. A. Bradford, E. P. Brinkman, T. F. J. van de Voorde, and C. Veen. 2016. Where, when and how plant–soil feedback matters in a changing world. Functional Ecology 30:1109–1121.

Wakano, J. Y. and L. Lehmann. 2014. Evolutionary branching in deme-structured populations. Journal of Theoretical Biology 351:83 – 95.

Wakano, J. Y., M. A. Nowak, and C. Hauert. 2009. Spatial dynamics of ecological public goods. Proceedings of the National Academy of Sciences 106:7910–7914.

Weiss, G. H. and M. Kimura. 1965. A mathematical analysis of the stepping stone model of genetic correlation. Journal of Applied Probability 2:129–149.

Weitz, J. S., C. Eksin, K. Paarporn, S. P. Brown, and W. C. Ratcliff. 2016. An oscillating tragedy of the commons in replicator dynamics with game-environment feedback. Proceedings of the National Academy of Sciences 113:E7518–E7525.

Wright, S. 1931. Evolution in mendelian populations. Genetics 16:97.

Wright, S. 1948. On the roles of directed and random changes in gene frequency in the genetics of populations. Evolution pp. 279–294.

